# Phagocytosis and self-destruction execute dendrite degeneration of *Drosophila* sensory neurons at distinct levels of NAD^+^ reduction

**DOI:** 10.1101/2020.06.26.173245

**Authors:** Hui Ji, Maria L. Sapar, Ankita Sarkar, Bei Wang, Chun Han

**Author notes:** These authors contributed equally to this work (Hui Ji, Maria L. Sapar).

## Abstract

After injury, severed dendrites and axons expose the “eat-me” signal phosphatidylserine (PS) on their surface and degenerate by disassembly. While axon degeneration is controlled by a conserved “axon-death” pathway that is thought to activate self-destruction, how PS exposure is regulated by this pathway and whether PS-induced phagocytosis contributes to neurite breakdown *in vivo* remain unknown. Here we show that in *Drosophila* sensory dendrites, PS exposure and self-destruction are triggered by two distinct levels of NAD^+^ reduction downstream of Sarm activation. Surprisingly, phagocytosis is the main driver of dendrite degeneration induced by both genetic NAD^+^ disruptions and injury. Furthermore, the axon-death factor Axed is only partially required for self-destruction of injured dendrites, acting in parallel with PS-induced phagocytosis. Lastly, injured dendrites exhibit a unique rhythmic calcium flashing that correlates with self-destruction. Therefore, a special genetic program coordinates PS exposure and self-destruction in injury-induced dendrite degeneration *in vivo*.

## INTRODUCTION

Physical insults to the nervous system often disrupt neuronal connectivity and function by damaging dendritic or axonal processes of neurons. Injured axons break down through a series of stereotypical events collectively called Wallerian degeneration (WD) (Coleman and Freeman, 2010; Waller, 1850). Dendrites undergo a similar program of degeneration after injury (Tao and Rolls, 2011). Before neurons can regenerate their processes and restore connections, the debris from damaged neurites has to be promptly cleared by phagocytes, which are cells that engulf dead cells or cell debris (Sapar and Han, 2019). Inefficient clearance can lead to neuroinflammation and further exacerbate the damage to the surrounding tissues (Davies et al., 2019; Galloway et al., 2019). Although WD is mainly considered to be a neurite-intrinsic, self-destructive process (Gerdts et al., 2016), whether phagocytosis actively contributes to degeneration of injured neurites *in vivo*, rather than merely passively removing neuronal debris, remains unclear.

WD is governed by an evolutionarily conserved pathway, which is also called “axon-death” pathway because it was discovered in studies focused primarily on axon degeneration in *Drosophila* and rodents (Freeman et al., 2003; Gerdts et al., 2016; Sapar and Han, 2019). This pathway is centered on nicotinamide adenine dinucleotide (NAD^+^), an essential cellular metabolite that is locally depleted after axons are injured. The signaling cascade starts with injury-induced activation of the E3 ubiquitin ligase Highwire/Phr1 in severed axons (Babetto et al., 2013; Xiong et al., 2012), which in turn causes degradation of nicotinamide mononucleotide adenyltransferase (Nmnat), an enzyme required for the synthesis of NAD^+^ (Zhai et al., 2009). The resultant decrease of NAD^+^ levels (Sasaki et al., 2016), together with an accumulation of the NAD^+^ precursor nicotinamide mononucleotide (NMN) (Di Stefano et al., 2015; Liu et al., 2018), activates Sarm/SARM1, a sterile alpha/Armadillo/Toll-Interleukin receptor homology domain protein (Bratkowski et al., 2020; Figley et al., 2021; Jiang et al., 2020; Osterloh et al., 2012; Shen et al., 2021). Sarm/SARM1 carries NADase activity; thus its activation causes catastrophic NAD^+^ depletion, which drives axon breakdown through unclear mechanisms (Gerdts et al., 2015; Wang et al., 2005).

Besides this core Highwire/Phr1-Nmnat-NAD^+^-Sarm/SARM1 pathway, several other factors have also been found to act in WD. Downstream of *Sarm, axundead* (*axed*) is required for axon degeneration of olfactory receptor neurons (ORNs) and wing sensory neurons in *Drosophila* (Neukomm et al., 2017). The loss of *axed* blocks axon degeneration even when Sarm is dominantly activated, raising the possibility that Axed activation, rather than NAD^+^ depletion, is the key switch of WD (Neukomm et al., 2017). In addition,*pebbled* (*peb*) encodes a *Drosophila* transcription factor required for axon degeneration of glutamatergic but not cholinergic sensory neurons in the wing (Farley et al., 2018). Lastly, the dual leucine kinase (DLK)/c-Jun N-terminal kinase (JNK) stress pathway also contributes to, but is not required for, WD in both *Drosophila* and mice (Miller et al., 2009; Yang et al., 2015). Although how these factors interact with the core pathway to promote axon degeneration is still mysterious, it is generally believed that catastrophic NAD^+^ depletion caused by Sarm activity initiates a neurite-intrinsic self-destruction program that ultimately is responsible for WD of axons (Babetto et al., 2013; Gerdts et al., 2015; Gerdts et al., 2016; Neukomm et al., 2017). While the WD pathway is primarily characterized in axons, evidence suggests that NAD^+^ reduction is also an essential step in injury-induced dendrite degeneration (Sapar et al., 2018; Tao and Rolls, 2011). However, which components of the WD pathway are conserved in dendrites remains unknown.

Neuronal debris is recognized by resident phagocytes of the nervous system through specific “eat-me” signals exposed on the neuronal surface. A highly conserved eat-me signal is phosphatidylserine (PS), a negatively charged phospholipid normally found in the inner leaflet of the plasma membrane of healthy cells. During apoptosis, PS is externalized to the outer leaflet of the plasma membrane to mark the cell for engulfment (Leventis and Grinstein, 2010). Genetic analyses of certain PS-binding bridging molecules and cell membrane receptors in mice and zebrafish suggest that PS recognition contributes to the phagocytosis of neurons (Fourgeaud et al., 2016; Mazaheri et al., 2014; Nandrot et al., 2007). Similarly, clearance of injured axons and dendrites in *Drosophila* requires Draper (Drpr), an engulfment receptor that binds to PS (Freeman et al., 2003; Han et al., 2014; MacDonald et al., 2006; Tung et al., 2013). PS exposure has also been directly observed on injured axons of mouse neurons *in vitro* (Shacham-Silverberg et al., 2018; Sievers et al., 2003) and on injured dendrites of *Drosophila* sensory neurons *in vivo* (Sapar et al., 2018). Although axonal PS exposure can be induced independent of axon degeneration *in vitro* (Shacham-Silverberg et al., 2018), ectopically induced PS exposure resulted in engulfment-dependent neurite reduction of otherwise healthy neurons in both the central nervous system (CNS) and the peripheral nervous system (PNS) of *Drosophila* (Sapar et al., 2018), pointing to a dominant effect of PS exposure in induced phagocytosis. Recent studies revealed a link between neuronal PS exposure and the WD pathway. Overexpression of Wld^S^, a fusion protein containing the full-length murine Nmnat1 (Mack et al., 2001), in *Drosophila* sensory neurons suppresses PS exposure of injured dendrites (Sapar et al., 2018). In addition, *Sarm1* ablation and NAD^+^ supplementation in neuronal culture reduce PS exposure on injured axons (Shacham-Silverberg et al., 2018). These observations raise two important questions: How does the WD pathway regulate and coordinate both neurite PS exposure and self-destruction? What are the relative contributions of PS-mediated phagocytosis and neurite self-destruction in WD *in vivo*?

To address these questions, we utilized *Drosophila* class IV dendritic arborization (C4da) neurons on the larval body wall, an established *in vivo* model of injury-induced dendrite degeneration (Han et al., 2014). In this system, degenerating dendrites of C4da neurons are phagocytosed by epidermal cells through the engulfment receptor Drpr. Here we show that PS exposure and dendrite self-destruction are two steps activated at different levels of NAD^+^reduction, and that *in vivo*, phagocytosis is the main driving force of dendrite degeneration induced by both genetic NAD^+^ perturbations and injury. Interestingly, injury and genetic NAD^+^perturbations cause dendrite degeneration through different cellular mechanisms in membrane disruption, dendrite calcium dynamics, and phagocytosis dependence. Furthermore, Axed specifically promotes, but is not required for, dendrite self-destruction, suggesting different mechanisms of injury-induced degeneration in axons and dendrites. Lastly, we report a previously unknown rhythmic calcium flashing specifically in injured dendrites, which is disrupted by Wld^S^ overexpression and *axed* loss, indicating a potential role of calcium flashing in dendrite self-destruction.

## RESULTS

### *Nmnat* knockout induces spontaneous dendrite degeneration due to Sarm-mediated NAD^+^reduction

NAD^+^ reduction is required for PS exposure on injured dendrites (Sapar et al., 2018). To determine if NAD^+^ loss is also sufficient to cause neuronal PS exposure, we decided to first investigate the impact of removing *Nmnat* from C4da neurons, as the loss of *Nmnat* is expected to cause cell-autonomous NAD^+^ reduction. *Nmnat* LOF is known to induce degeneration of eye photoreceptors (Zhai et al., 2006) and wing sensory neurons (Neukomm et al., 2017). However, a previous study found that C4da neurons mutant for *Nmnat* did not show dendrite degeneration, despite displaying dendrite reduction and axon degeneration (Wen et al., 2011). To reexamine the LOF phenotype of *Nmnat*, we used CRISPR-TRiM, a tissue-specific mutagenesis method we previously developed (Poe et al., 2019), to knock out *Nmnat* in C4da neurons. In this method, *Nmnat* is knocked out by C4da-specific *ppk-Cas9* and two ubiquitously expressed guide-RNAs (gRNAs) targeting *Nmnat*. To distinguish dendrite reduction caused by degeneration from that caused by growth defects, we used CD4-tdTomato (CD4-tdTom) to label C4da dendrites. Because tdTom is stable in phagosomes, the presence of tdTom-labeled dendrite debris in epidermal cells is an indication of dendrite breakdown and subsequent engulfment (Han et al., 2014).

As expected, *Nmnat* knockout (KO) in C4da neurons (Figure 1B) caused strong dendritic reduction (Figure 1G) in wandering 3^rd^ instar larvae as compared to the control (Figures 1A). In addition, dendrite debris was observed to spread in epidermal cells underneath and near the dendrites of *Nmnat* KO neurons (Figures 1B and 1H), suggesting that some dendrites had degenerated and were engulfed by epidermal cells. Unlike degenerating dendrites observed during developmental pruning or after injury (Han et al., 2011; Han et al., 2014), the remaining dendrites of *Nmnat* KO neurons did not show obvious blebbing or fragmentation, which may explain why dendrite degeneration was not detected previously in *Nmnat* mutant C4da neurons, considering that the membrane GFP marker used to label dendrites in the previous study is rapidly degraded by epidermal cells once engulfed and thus cannot visualize phagosomes (Han et al., 2014; Wen et al., 2011).

**Figure 1:**
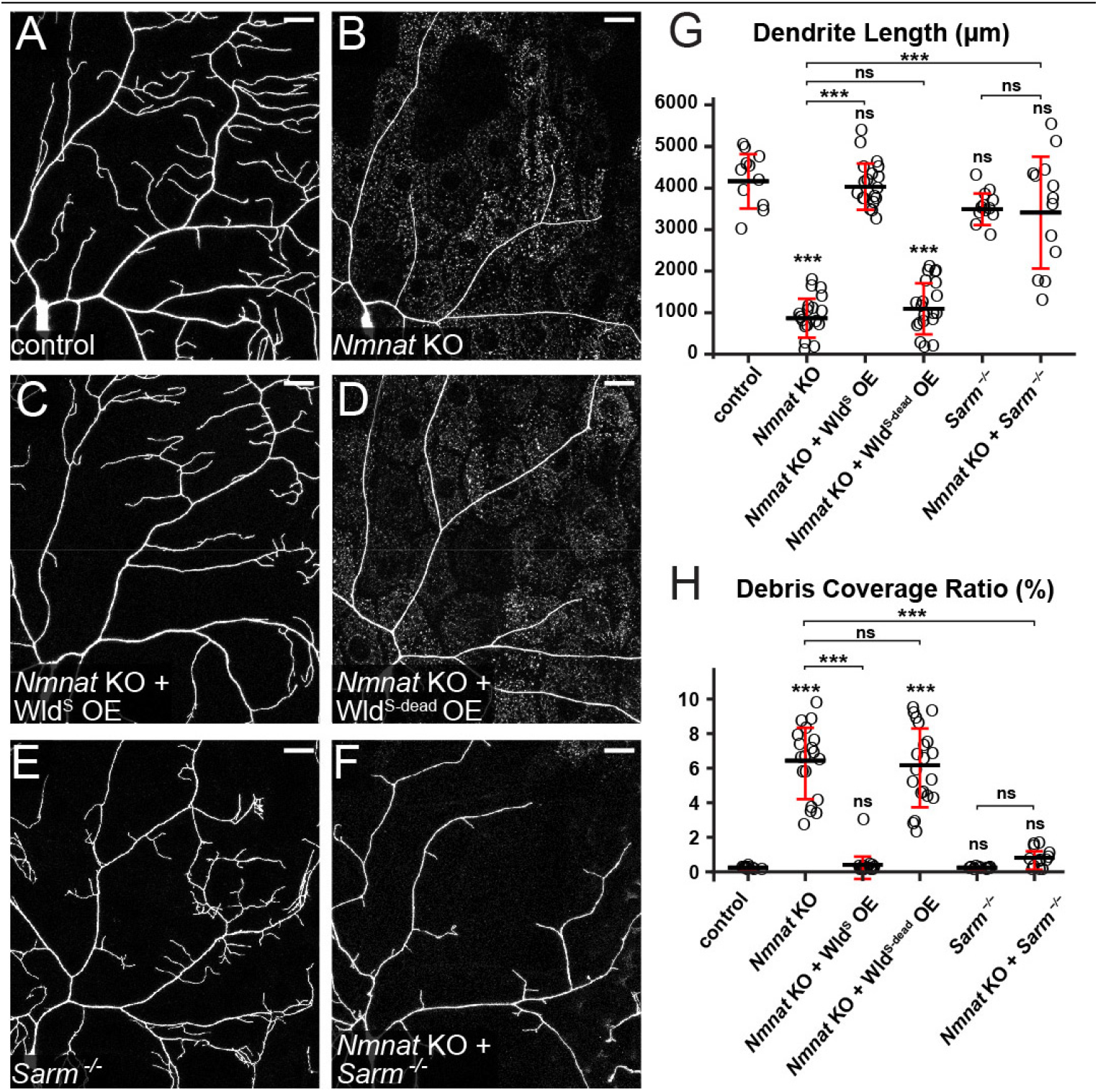
*Nmnat* knockout induces spontaneous dendrite degeneration due to Sarm-mediated NAD^+^ reduction. (A-F) Partial dendritic fields of control (A), *Nmnat* KO (B), *Nmnat* KO + Wld^S^ OE (C), *Nmnat* KO + Wld^S-dead^ OE (D), *Sarm^4705/4621^* (E), and *Nmnat* KO +*Sarm^4705/4621^* (F) ddaC C4da neurons. Neurons were labeled by *ppk>CD4-tdTom* (A-D) and *ppk-CD4-tdTom* (E and F). The C4da-specific KO was induced by *ppk-Cas9*. Scale bars, 25 μm. (G) Quantification of dendrite length. n = number of neurons: control (n =11, 6 animals); *Nmnat* KO (n = 19, 10 animals); *Nmnat* KO + Wld^S^ OE (n = 17, 10 animals); *Nmnat* KO + Wld^S-dead^ OE (n = 20, 11 animals); *Sarm^4705/4621^* (n = 12, 6 animals); *Nmnat* KO +*Sarm^4705/4621^* (n = 13, 8 animals). One-way ANOVA and Tukey’s test. (H) Quantification of debris coverage ratio, which is the percentage of debris area normalized by dendrite area ratio. Number of neurons: same as in (G). Kruskal-Wallis (One-way ANOVA on ranks) and Dunn’s test, p-values adjusted with the Benjamini-Hochberg method. For all quantifications, ***p≤0.001; n.s., not significant. The significance level above each genotype is for comparison with the control. Black bar, mean; red bar, SD.

Nmnat protects neurons by both synthesizing NAD^+^ and functioning as a chaperon protein (Ali et al., 2013). To verify that the observed dendrite degeneration is due to the loss of Nmnat enzymatic activity, we tried to rescue *Nmnat* KO neurons by overexpressing Wld^S^, which contains full NMNAT activity (Mack et al., 2001), and Wld^S-dead^, a mutant version of Wld^S^ that cannot synthesize NAD^+^ but maintains the chaperon function (Avery et al., 2009). Wld^S^ overexpression (OE) rescued the degeneration of *Nmnat* KO neurons and restored dendrite morphology to the wildtype level (Figures 1C, 1G, and 1H), while Wld^S-dead^ OE did not change dendrite length or debris level of *Nmnat* KO neurons (Figures 1D, 1G, and 1H). These results suggest that NAD^+^ reduction is responsible for the dendrite degeneration of *Nmnat* KO neurons.

We next asked whether Sarm plays a role in *Nmnat* KO-induced dendrite degeneration. Indeed, *Sarm* LOF completely blocked dendrite degeneration of *Nmnat* KO neurons (Figures 1F, 1G, 1H), as evident in the absence of dendrite debris in epidermal cells. However, these neurons showed more variable dendrite length (Figure 1G) compared to *Sarm* LOF alone (Figures 1E, 1G, and 1H), perhaps due to the loss of Nmnat chaperon function (Wen et al., 2011). Importantly, these results demonstrate that *Sarm* is required for the dendrite degeneration of *Nmnat* KO neurons. All the results above together suggest that *Nmnat* LOF in neurons cause spontaneous dendrite degeneration through Sarm-mediated NAD^+^ loss.

### PS exposure-mediated phagocytosis causes dendrite degeneration of *Nmnat* KO neurons

To ask whether *Nmnat* KO also causes neuronal PS exposure, we used an established method to visualize PS exposure on C4da dendrites. In this method, the fluorescent PS sensor GFP-Lact is expressed by the larval fat body and secreted into the hemolymph (Sapar et al., 2018). As C4da dendrites are largely exposed to the hemolymph, GFP-Lact labeling allows visualization of dynamic PS exposure on dendrites in intact live animals (Sapar et al., 2018). We found that GFP-Lact strongly labeled *Nmnat* KO neurons at distal branches that underwent degeneration (Figure 2B), while wildtype C4da neurons showed no labeling (Figure 2A). Interestingly, we also observed weaker GFP-Lact labeling on dendrite segments that did not display obvious signs of degeneration (outlined in Figure 2B), suggesting that PS exposure may precede dendrite breakdown, instead of being merely a consequence of dendrite degeneration. This conclusion was further corroborated by time-lapse imaging of *Nmnat* KO neurons (Video 1) and kymograph analysis (Figure 2C): PS exposure (indicated by white dotted lines) occurred well ahead of dendrite fragmentation (indicated by blue dotted lines).

**Figure 2:**
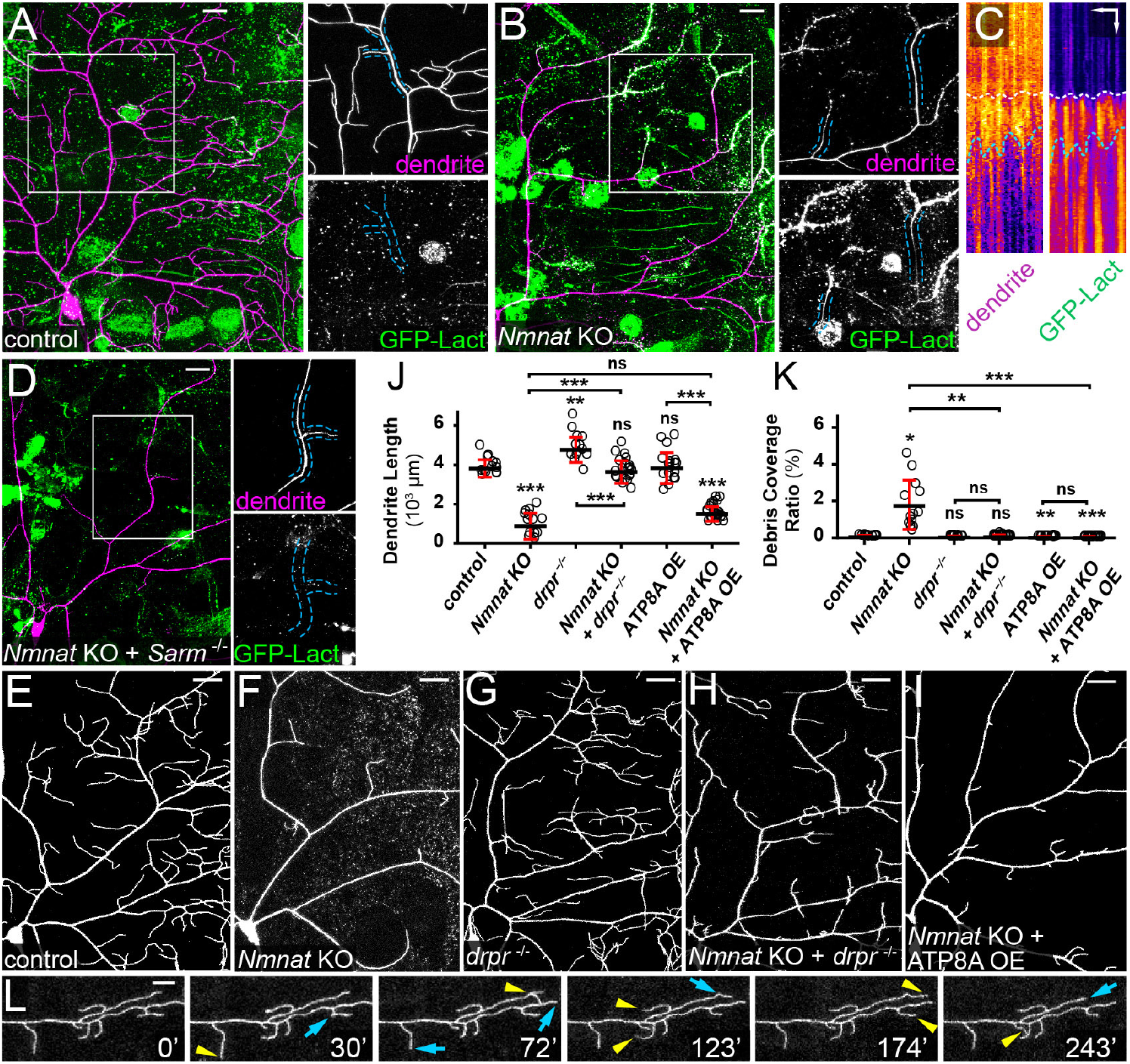
PS exposure-mediated phagocytosis causes dendrite degeneration of *Nmnat* KO neurons. (A) A control ddaC neuron showing the lack of GFP-Lact labeling. Insets show close-ups of an intact dendrite. Scale bar, 25 μm. (B) A *Nmnat* KO ddaC neuron undergoing spontaneous degeneration and showing labeling by GFP-Lact. In the inset, the outlined dendrite segments do not show obvious blebbing or fragmentation but are weakly labeled by GFP-Lact. Scale bar, 25 μm. (C) Kymographs of tdTom signal (dendrite) and GFP-Lact signal along a dendrite segment of a *Nmnat* KO ddaC neuron. White dotted lines indicate the timing of GFP-Lact initial binding to the dendrite; blue dotted lines indicate timing of the dendrite fragmentation based on the continuity of the dendrite signal. Scale bars, 10 μm horizontal, 60 min vertical. See also Video 1. (D) Dendrites of a *Nmnat* KO +*Sarm^4705/4621^* ddaC neuron showing the lack of GFP-Lact labeling. Insets show close-ups of an intact dendrite. Scale bar, 25 μm. (E-I) Partial dendritic fields of control (E), *Nmnat* KO (F), *drpr*^-/-^ (G), *Nmnat* KO +*drpr*^-/-^ (H) and *Nmnat* KO + ATP8A OE (I) neurons. Scale bars, 25 μm. (J) Quantification of dendrite length. n = number of neurons: control (n = 13, 7 animals); *Nmnat* KO (n = 13, 7 animals); *drpr*^-/-^ (n = 14, 8 animals); *Nmnat* KO +*drpr*^-/-^ (n = 22, 14 animals); ATP8A OE (n=17, 9 animals); *Nmnat* KO + ATP8A OE (n=23, 9 animals). One-way ANOVA and Tukey’s test. (K) Quantification of debris coverage ratio. Number of neurons: same as in (J). Kruskal-Wallis (One-way ANOVA on ranks) and Dunn’s test, p-values adjusted with the Benjamini-Hochberg method. (L) A time series of *Nmnat* KO +*drpr*^-/-^ dendrites. Yellow arrowheads indicate growth of dendrites compared to the previous frame; blue arrows indicate retractions of dendrites compared to the previous frame. Scale bar, 10 μm. See also Video 2. In all panels, neurons were labeled by *ppk-CD4-tdTom* (A-H) and *ppk-MApHS* (I). C4da-specific KO was induced by *ppk-Cas9*. For all quantifications, *p≤0.05, **p≤0.01, ***p≤0.001; n.s., not significant. The significance level above each genotype is for comparison with the control. Black bar, mean; red bar, SD.

As *Sarm* is required for dendrite degeneration of *Nmnat* KO neurons, we asked if Sarm also regulates PS exposure in these neurons. *Nmnat* KO neurons showed no PS exposure in the *Sarm* mutant background (Figure 2D), suggesting that Sarm-mediated NAD^+^ reduction is also responsible for inducing the observed PS exposure.

The observation of PS exposure on *Nmnat* KO dendrites raises the question of whether PS exposure contributes to dendrite breakdown by inducing phagocytic attacks from epidermal cells. Because PS-mediated epidermal engulfment of dendrites requires Drpr (Sapar et al., 2018), we examined *Nmnat* KO dendrites in *drpr* mutant larvae. Strikingly, *drpr* LOF completely blocked dendrite degeneration of *Nmnat* KO neurons (Figure 2H as compared to Figures 2E–2G; Figures 2J and 2K). These dendrites exhibited dynamic extension and retraction behaviors (Figure 2L and Video 2), demonstrating that they were not fragmented dendrites that failed to be cleared. Next, to examine whether dendrite breakdown induced by *Nmnat* KO depends on PS exposure, we tried to rescue *Nmnat* KO neurons by overexpressing ATP8A. ATP8A is the *Drosophila* ortholog of PS flippase responsible for keeping PS in the inner leaflet of the plasma membrane (Raiders et al., 2021; Tanaka et al., 2011), whose LOF in da neurons causes dendritic PS exposure (Sapar et al., 2018). Indeed, ATP8A OE completely blocked dendrite degeneration of *Nmnat* KO neurons and resulted in even less dendrite debris than the wildtype control (Figures 1I and 1K).

Thus, these data demonstrate that Sarm-mediated NAD^+^ reduction causes dendrite PS exposure, which in turn induces phagocytic attack to drive dendrite degeneration of *Nmnat* KO neurons.

### Wld^S^ OE and *Sarm* KO suppress PS exposure and fragmentation of injured dendrites

Wld^S^ protects injured neurites from degenerating by maintaining the NAD^+^ level in the neurites. We previously found that overexpressing Wld^S^ in C4da neurons blocked fragmentation and PS exposure of ablated dendrites at 10 hrs after injury (AI) (Sapar et al., 2018). To further investigate the role of the WD pathway in neuronal PS exposure, we examined the effects of overexpressing Wld^S^ and knocking out *Sarm* in C4da neurons at 24 hrs after laser-severing of dendrites. Neuronal-specific KO was conducted using *SOP-Cas9*, which is active in precursor cells of da neurons, to minimize potential gene perdurance (Poe et al., 2019). As expected, Wld^S^ OE blocked degeneration and clearance of injured dendrites also at 24 hrs AI (Figure 3C, as compared to the control in Figure 3B), even though the injured arbors were greatly simplified as compared to uninjured dendrites (Figures 3A and 3E). Neuronal-specific *Sarm* KO showed similar effects in injured dendrites (Figures 3D and 3E) to those of Wld^S^ OE.

**Figure 3:**
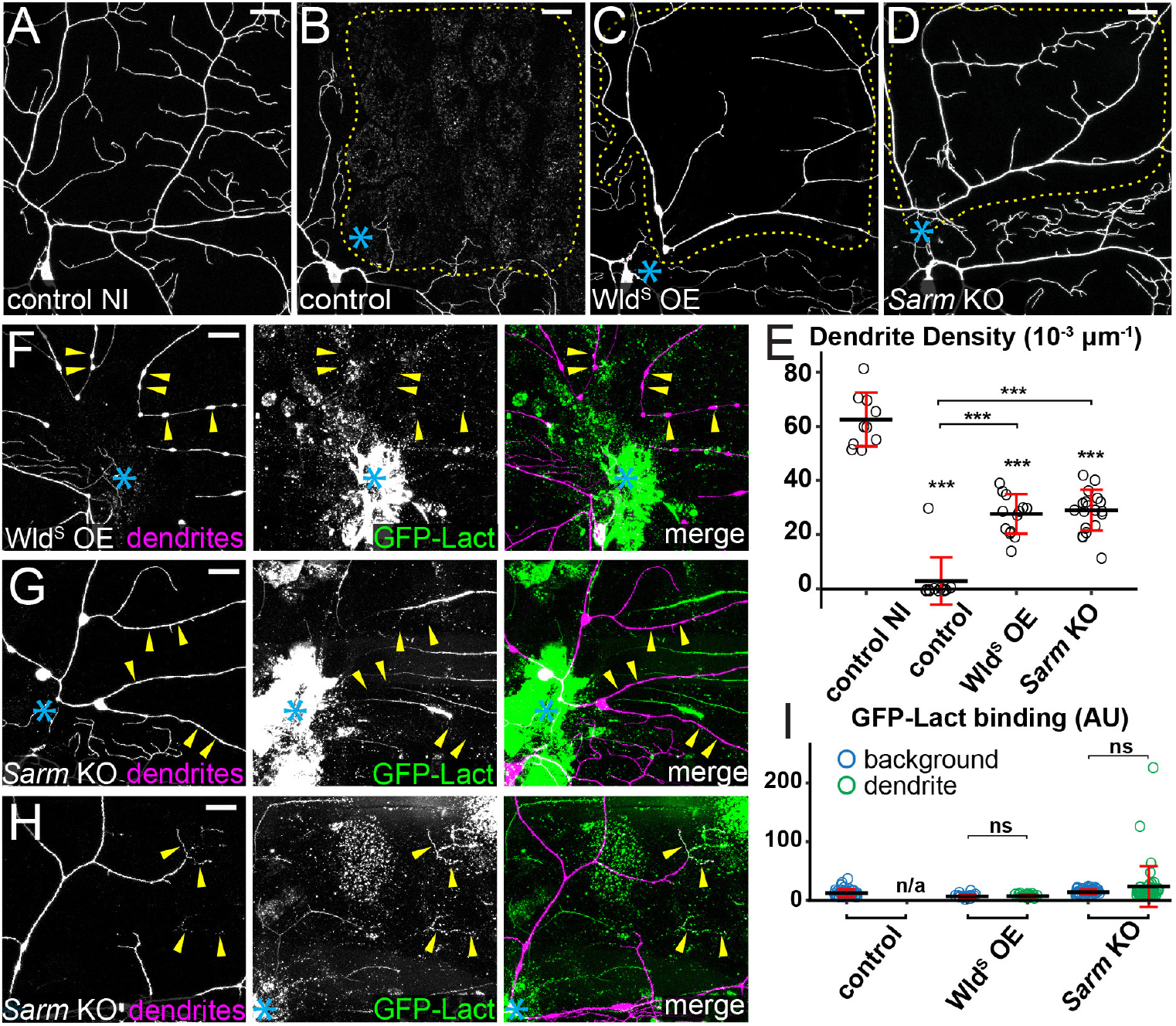
Wld^S^ OE and *Sarm* KO suppress PS exposure and fragmentation of injured dendrites. (A) Partial dendritic field of an uninjured ddaC control neuron. (B-D) Partial dendritic fields of control (B), Wld^S^ OE (C) and *Sarm* KO (D) ddaC neurons 24 hrs after injury (AI). Blue asterisk, injury site. Yellow dots outline regions covered by injured dendrites. (E) Quantification of dendrite density (1000 x dendritic length/area) in regions covered by injured dendrites. n = number of neurons: control no injury (NI) (n = 10, 10 animals); control (n = 12, 6 animals); Wld^S^ OE (n = 13, 6 animals); *Sarm* KO (n = 19, 6 animals). (F and G) Wld^S^ OE (F) and *Sarm* KO (G) ddaC neurons 24 hrs AI, showing unfragmented dendrites and lack of GFP-Lact binding to severed dendrites (yellow arrowheads). (H) Dendrites of a *Sarm* KO ddaC neuron 24 hrs AI, showing infrequent GFP-Lact binding at fragmented terminal branches (yellow arrowheads). (I) Quantification of GFP-Lact binding on injured dendrites. The GFP-Lact intensity is shown for both background epidermal regions and injured dendrites. The outliers in *Sarm* KO dendrite dataset correspond to infrequent degenerating terminal branches. Background: epidermal regions without dendrites. n = number of measurements: control background (n = 39) and control dendrite (n/a, no remaining dendrites), 7 animals; Wld^S^ OE background (n = 18) and Wld^S^ OE dendrite (n = 18), 4 animals; *Sarm* KO background (n = 48) and *Sarm* KO dendrite (n = 48), 9 animals. In all panels, neurons were labeled by *ppk-MApHS* (A-C; only tdTom channel is shown), *ppk>CD4-tdTom* (D), and *ppk-CD4-tdTom* (F-H). Neuronal-specific KO was induced by *SOP-Cas9*. For all quantifications, One-way ANOVA and Tukey test; ***p≤0.001; n.s., not significant; n/a., not applicable. The significance level above each genotype in (E) is for comparison with the control. Black bar, mean; red bar, SD. Scale bars, 25 μm.

We next examined the effects of Wld^S^ OE and *Sarm* KO on dendritic PS exposure 24 hrs AI. Wld^S^ OE prevented PS exposure on severed dendrites, even on branches that showed blebbing and breakage (Figures 3F and 3I). *Sarm* KO also effectively blocked PS exposure on injured dendrites (Figures 3G and 3I), with occasionally strong PS labeling on fragmented distal terminal branches (Figures 3H and 3I). The above data in dendrite injury together suggest that Sarm-mediated NAD^+^ reduction causes both PS exposure and degeneration of injured dendrites. The PS exposure on fragmented terminal dendrites of *Sarm* KO neurons could be due to relatively faster NAD^+^ turnover in those local branches even in the absence of Sarm.

### Ectopic PS exposure causes injured dendrites of Wld^S^ OE and *Sarm* KO neurons to degenerate

The absence of PS exposure on injured dendrites of Wld^S^ OE and *Sarm* KO neurons raises the question of whether the degeneration defects of these neurons are due to the lack of PS-mediated epidermal attack on dendrites. To test this hypothesis, we ectopically induced neuronal PS exposure by knocking out *CDC50*, which encodes a chaperone necessary for the function of *Drosophila* flippases (Tanaka et al., 2011), and by overexpressing TMEM16F, a mammalian PS scramblase (Segawa et al., 2011). These manipulations in C4da neurons cause mild but appreciable phagocytosis-dependent dendrite loss (Sapar et al., 2018). *CDC50* KO or TMEM16F OE alone in Wld^S^ OE neurons caused partial or complete degeneration of injured dendrites and the accompanying epidermal engulfment (Figures 4A, 4B, and 4E). A much stronger effect was observed when *CDC50* KO and TMEM16F OE were combined in Wld^S^ OE neurons: Degeneration and clearance of injured dendrites were restored to the wildtype level (Figures 4C and 4E). Similarly, *CDC50* KO and TMEM16F OE caused injured dendrites of *Sarm* KO neurons to completely degenerate (Figures 4D and 4E). Because injury induces a more rapid and stronger PS exposure on dendrites than *CDC50* KO or TMEM16F OE (Sapar et al., 2018), our data strongly suggest that injury-induced PS exposure is a major driver of dendrite breakdown by activating phagocytic attack by epidermal cells.

**Figure 4:**
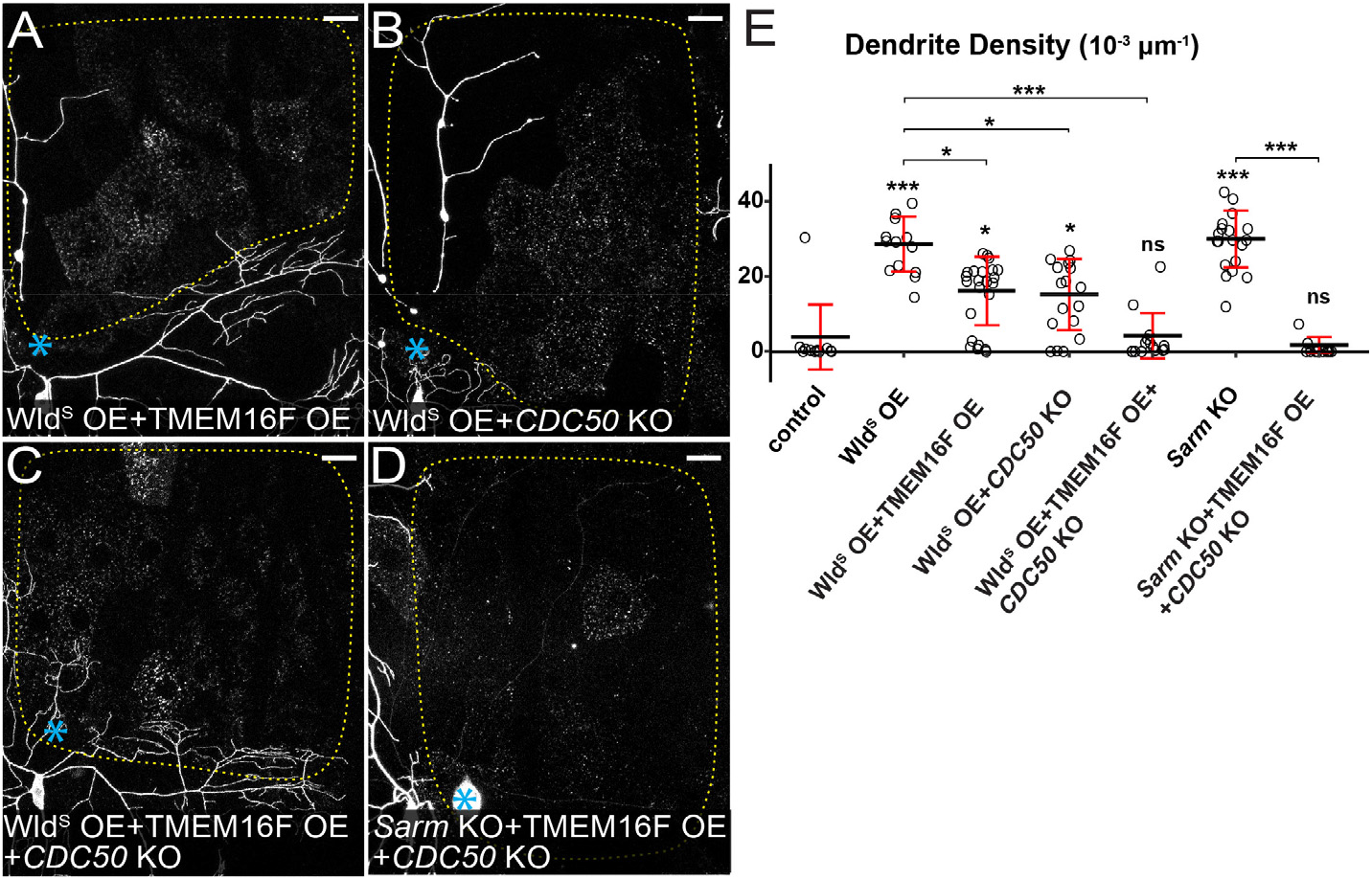
Ectopic PS exposure causes injured dendrites of Wld^S^ OE and *Sarm* KO neurons to degenerate. (A-D) Partial dendritic fields of Wld^S^ OE + TMEM16F OE (A), Wld^S^ OE + *CDC50* KO (B), Wld^S^ OE +TMEM16F OE + *CDC50* KO (C) and *Sarm* KO + TMEM16F OE + *CDC50* KO (D) ddaC neurons 24 hrs AI. Blue asterisk, injury site. Yellow dots outline regions covered by injured dendrites. Neurons were labeled by *ppk-MApHS* (A, B, and C) and *ppk-CD4-tdTom* (D). Neuronal-specific KO was induced by *ppk-Cas9* (B and C) and *SOP-Cas9* (D). Scale bars, 25 μm. (E) Quantification of dendrite density. n = number of neurons: control (n = 12, 6 animals); Wld^S^ OE (n = 13, 6 animals); Wld^S^ OE + TMEM16F OE (n = 23, 8 animals); Wld^S^ OE +*CDC50* KO (n = 17, 8 animals); Wld^S^ OE + TMEM16F OE + *CDC50* KO (n = 16, 8 animals); *Sarm* KO (n = 19, 6 animals); *Sarm* KO + TMEM16F OE +*CDC50* KO (n = 13, 7 animals). Kruskal-Wallis (One-way ANOVA on ranks) and Dunn’s test, p-values adjusted with the Benjamini-Hochberg method. The dataset for Wld^S^ OE and *Sarm* KO is the same as in Figure 3E. For all quantifications, ^*^p≤0.05, ^***^p≤0.001; n.s., not significant. The significance level above each genotype is for comparison with the control. Black bar, mean; red bar, SD.

### Injury and NAD^+^ depletion cause phagocytosis-independent dendrite self-destruction

Although PS-mediated phagocytosis is sufficient to break down injured dendrites, blocking phagocytosis with *drpr* LOF failed to prevent the fragmentation of injured dendrites 20 hrs AI, even though the dendrite debris was left unengulfed (Figures 5A–5C). A neurite self-destruction program triggered by NAD^+^ depletion is known to drive WD of cultured neurons in the absence of phagocytes (Gerdts et al., 2015). To test the possibility that a further NAD^+^ reduction beyond the level that triggers PS exposure causes phagocytosis-independent dendrite self-destruction, we tested the effect of overexpressing a gain-of-function Sarm (Sarm^GOF^) in C4da neurons, because Sarm^GOF^ potently depletes NAD^+^ and causes dominant neurodegeneration (Neukomm et al., 2017). Indeed, Sarm^GOF^ OE caused complete dendrite degeneration in most neurons as early as 24 hrs after egg laying (AEL) (Figures 5D and 5J), while *Nmnat* KO did not cause dendrite degeneration until 88-91 hrs AEL. Supporting the role of engulfment in NAD^+^ reduction-induced dendrite degeneration, *drpr* LOF strongly suppressed dendrite degeneration of Sarm^GOF^ OE neurons at both 24 hrs AEL and 48 hrs AEL (Figures 5E, 5G, and 5J). However, *drpr* LOF failed to prevent dendrite fragmentation of Sarm^GOF^ OE neurons at wandering 3^rd^ instar (Figures 5H–5J), suggesting that NAD^+^ depletion is able to cause phagocytosis-independent dendrite self-destruction. These results together support the hypothesis that dendrite self-destruction induced by NAD^+^ depletion is responsible for the fragmentation of injured dendrites when phagocytosis is suppressed.

**Figure 5:**
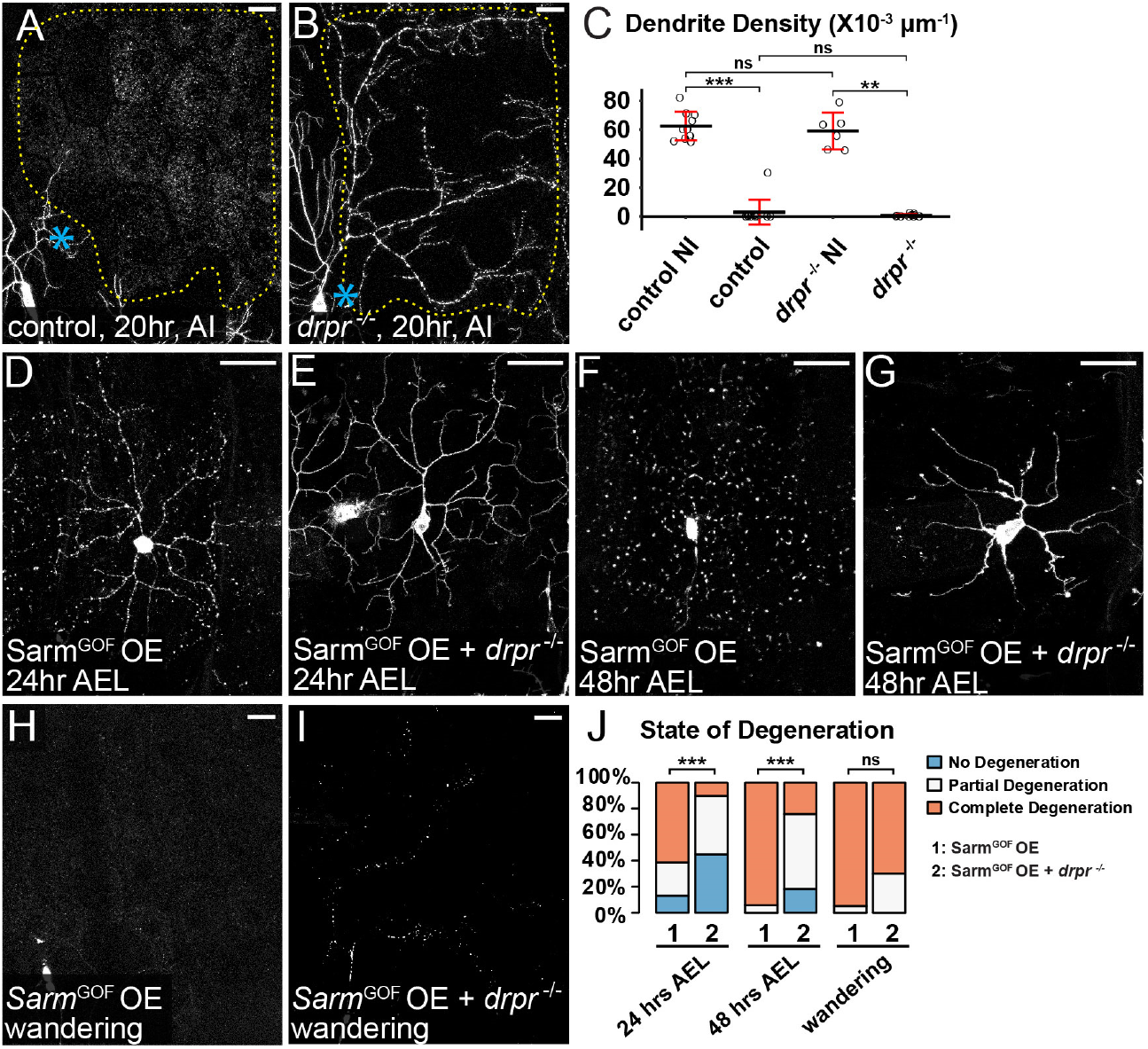
Injury and NAD^+^ depletion cause phagocytosisindependent dendrite self-destruction. (A-B) Partial dendritic fields of ddaC neurons in control (A) and *drpr*^-/-^ (B) animals 20 hrs AI. The dendrite debris lining up in the original dendritic pattern in (B) indicates dendrite fragmentation and the lack of epidermal engulfment. Blue asterisk, injury site. Yellow dots outline regions covered by injured dendrites. (C) Quantification of dendrite density. n = number of neurons: control no-injury (NI) (n = 10, 10 animals); control AI (n = 12, 6 animals); *drpr*^-/-^ NI (n = 6, 5 animals); *drpr*^-/-^ AI (n = 13, 5 animals). Kruskal-Wallis (One-way ANOVA on ranks) and Dunn’s test, p-values adjusted with the Benjamini-Hochberg method. (D-G) ddaC neurons in *Sarm*^GOF^ OE 24 hrs after egg laying (AEL) (D), *Sarm*^GOF^ OE +*drpr*^-/-^ 24 hrs AEL (E), *Sarm*^GOF^ OE 48 hrs AEL (F) and *Sarm*^GOF^ OE +*drpr*^-/-^ 48 hrs AEL (G). (H-I) Partial dendritic fields of *Sarm*^GOF^ OE (H) and *Sarm*^GOF^ OE + *drpr*^-/-^ (I) ddaC neurons at the wandering stage. (J) Quantification of dendrite degeneration showing percentages of neurons undergoing partial degeneration, complete degeneration, and no degeneration. n = number of neurons: *Sarm*^GOF^ OE 24 hrs AEL (n = 31, 7 animals); *Sarm*^GOF^ OE +*drpr*^-/-^ 24 hrs AEL (n = 49, 17 animals); *Sarm*^GOF^ OE 48 hrs AEL (n = 17, 7 animals); *Sarm*^GOF^ OE +*drpr*^-/-^ 48 hrs AEL (n = 33, 11 animals); *Sarm*^GOF^ OE wandering (n = 19, 14 animals); *Sarm*^GOF^ OE +*drpr*^-/-^ wandering (n = 10, 6 animals). Freeman-Halton extension of Fisher’s exact test. In image panels above, neurons were labeled by *ppk-MApHS* (A and B) and *ppk>CD4-tdTom* (D-I). C4da-specific KO was induced by *ppk-Cas9*. For all quantifications, **p≤0.01, ***p≤0.001; n.s., not significant. Black bar, mean; red bar, SD. Scale bars, 25 μm.

### Axed promotes self-destruction of injured dendrites in parallel with PS-mediated phagocytosis

To understand how other components of the WD pathway regulate dendrite degeneration, we examined LOF of *axed* and *peb*. Tissue-specific KO of *axed* in Or22a olfactory neurons, an established axon-injury model (MacDonald et al., 2006), effectively blocked degeneration of severed axons 7 days after antenna ablation (Figure S1A–S1D, and S1G), demonstrating the effectiveness of the gRNAs. In C4da neurons, both KO and knockdown (KD) of *axed* also efficiently blocked injury-induced axon degeneration (Figures S1H–S1K), consistent with the role of Axed in WD. To test if Axed is involved in PS exposure-induced dendrite degeneration resulting from *Nmnat* LOF, we knocked out both *Nmnat* and *axed* in da neurons. Unlike *Sarm* KO, which fully rescued *Nmnat* KO-induced degeneration (Figures S1M, S1O, S1P, and S1Q), *axed* KO did not affect dendrite degeneration caused by *Nmant* KO (Figures S1L, S1N, S1P, and S1Q), suggesting that *axed* is not required for the PS exposure induced by NAD^+^ reduction.

We next asked if Axed is involved in injury-induced dendrite degeneration. Injured dendrites of wildtype neurons completely fragmented by 10 hrs AI (Figures 6A and 6E). *axed* KO in neurons resulted in incomplete and no fragmentation of injured dendrites only in small fractions (25% and 8%, respectively) of neurons (Figures 6B and 6E), suggesting that *axed* is not absolutely required for the degeneration of injured dendrites. We next asked if Axed specifically contributes to self-destruction of injured dendrites. Because PS-induced phagocytosis may be sufficient to break down injured dendrites by 10 hrs AI, thereby masking dendrite protection by *axed* KO, we examined the effect of *axed* KO in *drpr* mutant. *drpr* LOF alone caused incomplete and no dendrite fragmentation in 36% and 23% injured neurons, respectively (Figures 6C and 6E), consistent with the idea that phagocytosis contributes significantly to dendrite breakdown at this stage. In comparison, *axed* KO in *drpr* mutant strongly blocked dendrite degeneration at 10 hrs AI, with 56% incomplete fragmentation and 36% no fragmentation (Figures 6D and 6E), suggesting that *axed* KO and *drpr* LOF have an additive effect in dendrite protection. Surprisingly, this combined protection was significantly weakened by 24 hrs AI, with only 47% incomplete fragmentation and 5% no fragmentation (Figures 6F–6J). These data suggest that Axed promotes dendrite self-destruction in parallel with PS-induced phagocytosis; however, other factors can also cause self-destruction in the absence of Axed.

**Figure 6:**
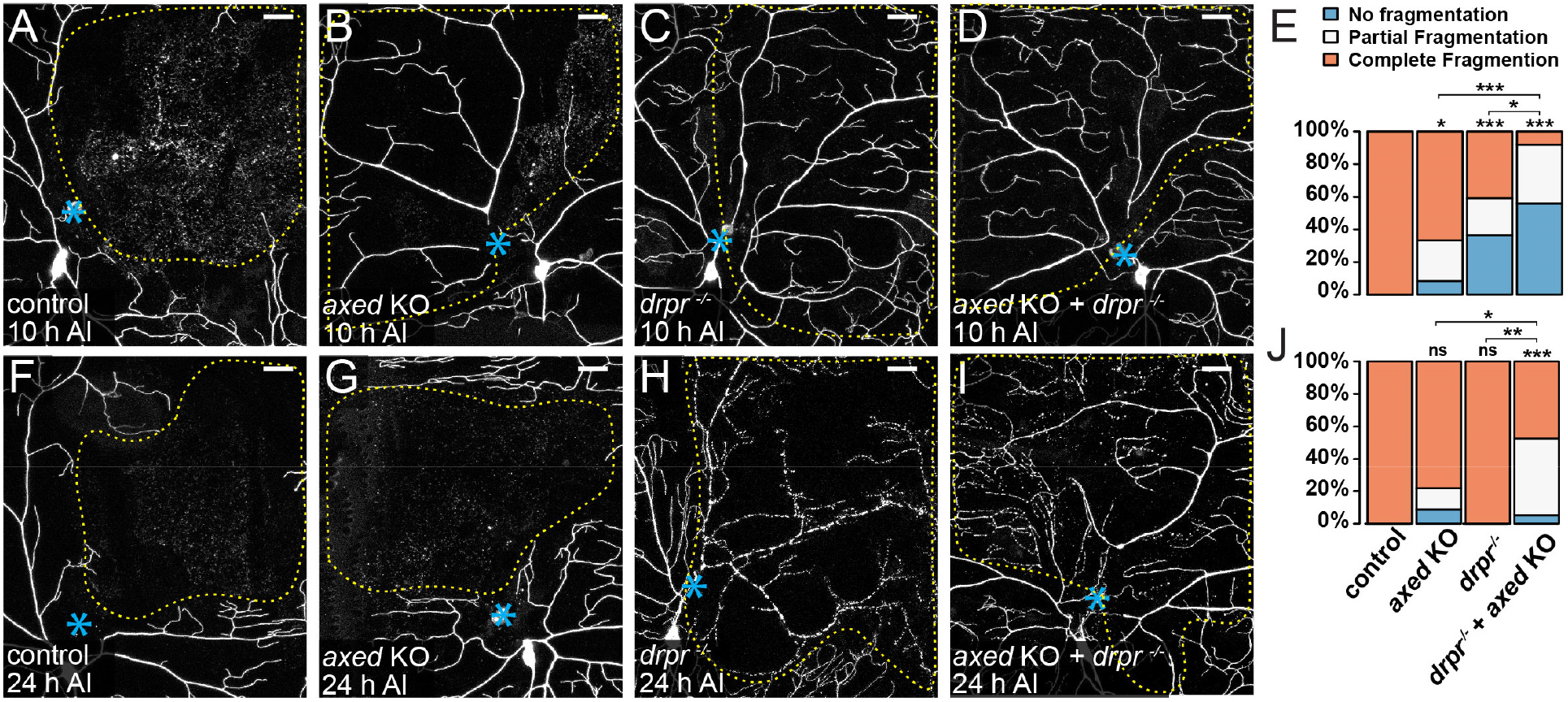
Axed drives dendrite self-destruction in parallel with PS-mediated phagocytosis. (A-D) Partial dendritic fields of wildtype (A), *axed* KO (B), *drpr*^-/-^ (C), and *axed* KO +*drpr*^-/-^ (D) ddaC neurons 10 hrs AI. (B) shows partial fragmentation; (C) and (D) show no fragmentation. (E) Quantification of dendrite fragmentation showing percentages of neurons undergoing no fragmentation, partial fragmentation, and complete fragmentation of injured dendrites at 10 hrs AI. n = number of neurons: wildtype (n=18, 9 animals); *axed* KO (n = 24, 10 animals); *drpr*^-/-^ (n = 22, 11 animals); *axed* KO +*drpr*^-/-^ (n = 25, 11 animals). Freeman-Halton extension of Fisher’s exact test. (F-I) Partial dendritic fields of wildtype (F), *axed* KO (G), *drpr*^-/-^ (H), and *axed* KO +*drpr*^-/-^ (I) ddaC neurons 24 hrs AI. (I) shows partial fragmentation. In all image panels, blue asterisks indicate injury sites; yellow dots outline regions covered by injured dendrites. Neurons were labeled by *ppk>CD4-tdTom* (A, B, F and G) and *ppk-CD4-tdTom* (C, D, H, and I). Neuronal-specific KO was induced by *SOP-Cas9* (B, D, G, and I). Scale bars, 25 μm. (J) Quantification of dendrite fragmentation showing percentages of neurons undergoing no fragmentation, partial fragmentation, and complete fragmentation of injured dendrites at 24 hrs AI. n =number of neurons: wildtype (n=21, 12 animals); *axed* KO (n = 23, 10 animals); *drpr*^-/-^ (n = 12, 5 animals); *axed* KO +*drpr*^-/-^ (n = 19, 8 animals). Freeman-Halton extension of Fisher’s exact test. For all quantifications, *p≤0.05, **p≤0.01, ***p≤0.001; n.s., not significant. The significance level above each genotype is for comparison with the wildtype.

In contrast to *axed, peb* KO in C4da neurons did not block dendrite fragmentation after injury but altered dendrite morphology (Figures S1R–S1T). Unexpectedly, knocking out *peb* from precursors of Or22a neurons caused loss of some or all of Or22a axons in uninjured adult brains (Figure S1G), suggesting that *peb* plays a role in Or22a development or axon patterning. The remaining axons of *peb* KO neurons did not show defects in axon degeneration after injury (Figures S1E–S1G). These results suggest that Peb may be a neuronal type-specific modulator of the WD pathway and is not required for dendrite degeneration of da neurons.

### Injured dendrites undergo severe membrane rupture during dendrite fragmentation

To further understand how injury induces dendrite degeneration, we investigated the extent of membrane rupture during dendrite breakdown using a split GFP-based assay. In this membrane rupture assay (Figure 7A), neurons express myristoylated tdTom-GFP(1-10) and the fat body secretes GFP(11)x7 into the hemolymph. GFP(11)x7 will bind myr-tdTom-GFP(1-10) and reconstitute fluorescent GFP only when the dendrite membrane is ruptured to allow diffusion of GFP(11)x7 into the cytoplasm of neurons. While reconstituted GFP was not detected in uninjured wildtype dendrites (Figure S2A) or degenerating dendrites of *Nmnat* KO neurons(Figures 7B, open arrowheads, and 7D–7E), it was observed in severed wildtype dendrites that were undergoing blebbing and fragmentation (Figures 7C, arrowheads, and 7E), as well as on membrane pieces shed from injured dendrites (Figure 7C, arrows, and 7D). In time-lapse movies, GFP signals were visible at low levels in injured branches soon after laser ablation, likely due to GFP(11)x7 entry through the injury site. The signals were kept at constantly low levels until injured dendrites fragmented, when GFP signals rapidly increased in large dendrite particles (Video 3). These results suggest that injured dendrites undergo severe membrane rupture during fragmentation. In contrast, *Nmnat* KO neurons experience much milder disruptions of membrane integrity, even though they are losing membranes due to the attack of epidermal cells, probably because their dynamically growing branches are efficient in repairing membrane damages.

**Figure 7.**
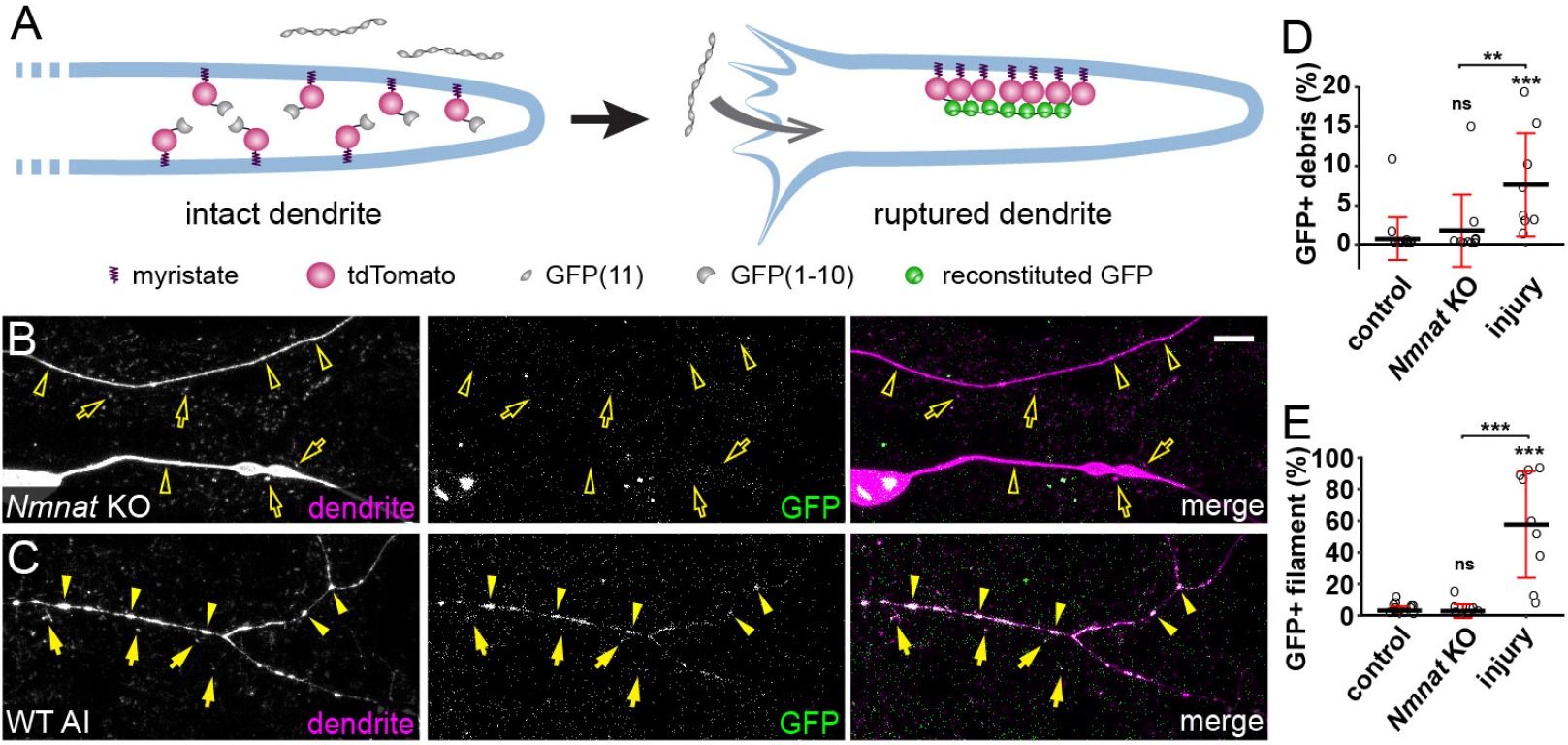
Injured dendrites undergo severe membrane rupture during dendrite fragmentation. (A) A diagram for the membrane rupture assay. Extracellular GFP(11)_x7_ is separated from intracellular myr-tdTom-GFP(1-10) attached to the inner membrane of dendrites. Fluorescent GFP is reconstituted only when the dendrite membrane is ruptured to allow diffusion of GFP(11)_x7_ into the cytoplasm of neurons. (B) *Nmnat* KO dendrites (open arrowheads) and debris (open arrows) lacking reconstituted GFP in the membrane rupture assay. Scale bar, 25 μm. (C) Reconstituted GFP labeling of injured wildtype dendrites (arrowheads) and debris (arrows) in the membrane rupture assay. Scale bar, 25 μm. See also Video 3. (D) Quantification of GFP-positive debris, showing the percentage of GFP-positive debris area in tdTom-positive debris area. Number of ROIs: wildtype NI (n= 15, 7 animals); *Nmnat* KO NI (n= 10, 6 animals) wildtype 6 hrs AI (n= 10, 4 animals). Kruskal-Wallis (One-way ANOVA on ranks) and Dunn’s test, p-values adjusted with the Benjamini-Hochberg method. (E) Quantification of GFP-positive filament, showing the percentage of GFP-positive filament area in tdTom-positive filament area. Number of ROIs: same as in (D). Kruskal-Wallis (One-way ANOVA on ranks) and Dunn’s test, p-values adjusted with the Benjamini-Hochberg method. For all quantifications, **p≤0.01; ***p≤0.001; n.s., not significant. The significance level above each genotype is for comparison with the control. Black bar, mean; red bar, SD.

### Injured dendrites exhibit calcium flashing that is suppressed by Wld^S^ and *axed* loss

The observed membrane rupture of injured dendrites is consistent with phagocytic attacks of epidermal cells on PS-exposing dendrites (Figure 7C). To understand potential signaling events that may lead to dendritic PS exposure and membrane rupture, we examined calcium dynamics in injured dendrites using long-term time-lapse imaging (Ji and Han, 2020; Poe et al., 2017), because calcium influx is essential for and correlates with degeneration of damaged axons in neuronal culture and *in vivo* (George et al., 1995; Vargas et al., 2015; Williams et al., 2014). A previous study in zebrafish identified an initial calcium influx at the time of axon severing and a second calcium wave that coincides with axon fragmentation (Vargas et al., 2015). We recorded calcium dynamics using a membrane-tethered GCaMP6s (myr-GCaMP6s) (Akbergenova et al., 2018), which detected only occasional local rise of calcium in dendrites of wildtype (Video 4) and *Nmnat* KO neurons (Video 5). In the injury model, we observed an initial calcium rise immediately after laser injury in both detached dendrites and the rest of the neurons (Figure 8A, 2s AI). Interestingly, soon after calcium dropped to the baseline level, severed dendrites but not those connected to the cell body entered a phase of continuous calcium flashes that lasted 1-5 hours (Figures 8A at 1h AI, 8D, and Video 6). Afterward, severed dendrites stayed relatively quiescent for 1-3 hrs before calcium surged again at the time of dendrite blebbing and fragmentation (Figures 8A at 6h AI, 8D, and Video 6). All severed dendrites but not those still attached to cell bodies showed flashing, quiescence, and surge phases after injury, even though the exact timing and duration of each phase varied from dendrite to dendrite (Figures S3A and S3D).

**Figure 8:**
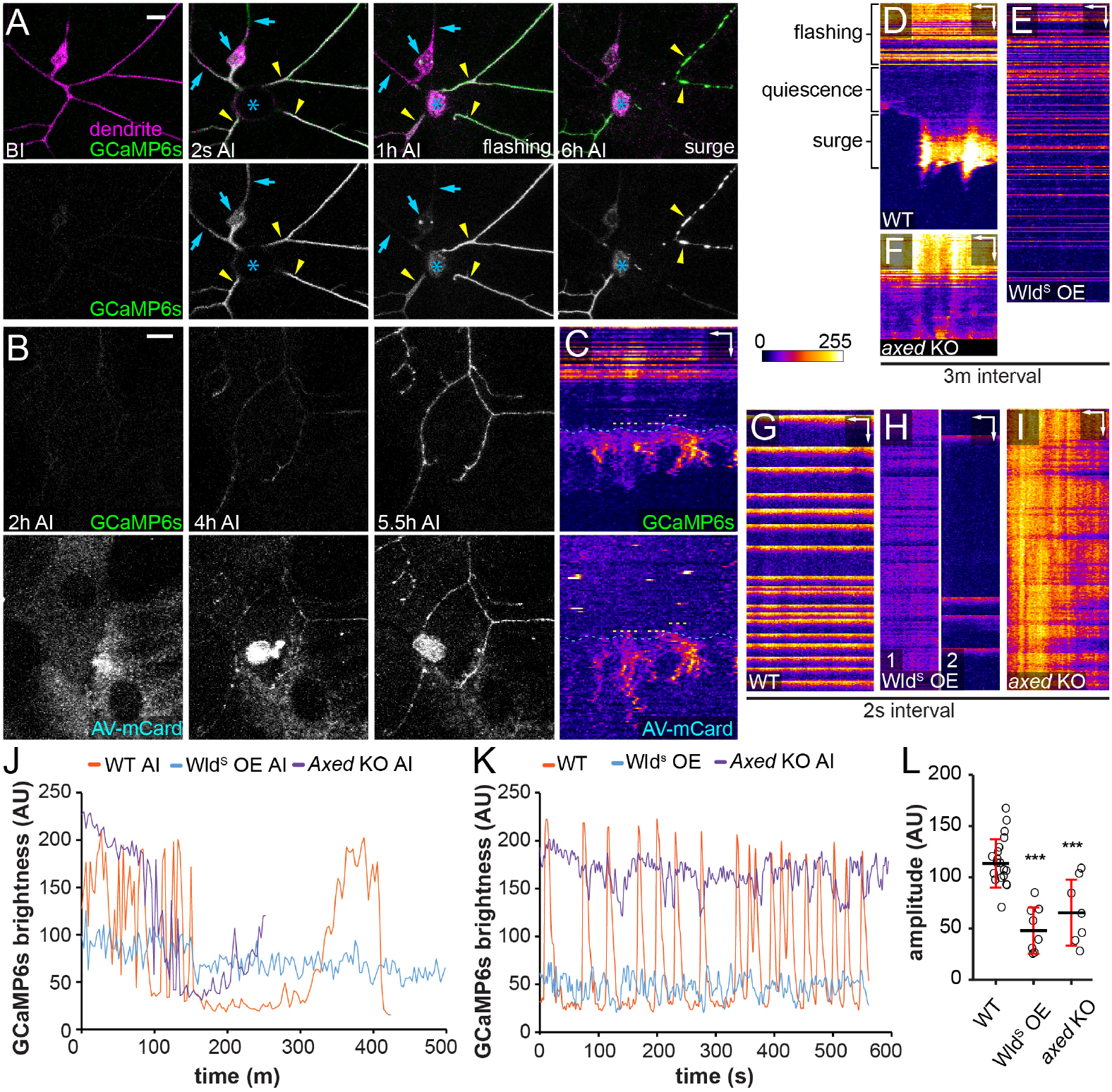
Calcium flashing precedes PS exposure and fragmentation of injured dendrites. (A) A ddaC neuron in a 3^rd^ instar larva expressing myr-GCaMP6s before (BI) and after (2s AI, 1h AI, and 6h AI) laser injury. Blue arrows, uninjured dendrites or cell body; yellow arrowheads, injured dendrites; blue asterisk, injury site. Scale bar, 10 μm. See also Video 6. (B) Labeling of injured ddaC dendrites expressing GCaMP6s by AV-mCard after (2 h, 4 h, and 5.5 h) injury. Scale bar, 10 μm. See also Video 7. (C) Kymograph of GCaMP6s signal and AV-mCard binding on an injured dendrite shown in B. Yellow dotted lines, timing of AV-mCard binding; blue dotted lines, timing of the final calcium surge. Scale bars, 3 μm horizontal, 40 min vertical. (D-F) Kymograph of GCaMP6s signal in wildtype (D), Wld^S^-overexpressing (E), and *axed* KO (F) dendrites after injury with 3-min interval. Scale bar, 5 μm horizontal, 60 min vertical. See also Videos 6, 8 and 12. (G-I) Kymograph of GCaMP6s signal in wildtype (G), Wld^S^-overexpressing (H), and *axed* KO (I) dendrites after injury with 2-s interval. See also Videos 9-11 and 13. Scale bars, 5 μm horizontal, 45 sec vertical. (J) Plot of GCaMP6s brightness in injured dendrites in (D), (E), and (F) over time with 3-m interval. Measurements in injured dendrites start from 0.5-1 hr AI. (K) Plot of GCaMP6s brightness in injured dendrite in (G), (H), and (I) over time with 2-s interval. Measurements start from 0.5-1 hr AI. (L) Quantification of average flashing amplitude (GCaMP6s brightness difference between maximum peak and neighboring minimum peak) captured in 2 s-interval time-lapse images. n=number of dendrite arbors: wildtype 0.5-1.5 hrs AI (n= 19, 5 animals); Wld^S^-overexpressing 2 hrs AI (n=8, 4 animals); *axed* KO 0.5-1 hrs AI (n=7, 3 animals). One-way ANOVA and Tukey’s test. For all quantifications, ***p≤0.001; n.s., not significant. The significance level above each genotype is for comparison with the control. Black bar, mean; red bar, SD.

It was reported that nanoscale ruptures of the axonal plasma membrane are responsible for extracellular calcium influx in neuroinflammatory lesions (Witte et al., 2019) and after spinal cord contusion (Williams et al., 2014). To determine whether the last calcium rise that coincides with dendrite degeneration could be a result of phagocytosis-induced membrane breakage, we imaged both calcium dynamics and PS exposure of injured dendrites using the PS sensor Annexin V-mCardinal (AV-mCard) (Sapar et al., 2018). AV-mCard labeling of dendrites appeared slightly ahead of the final calcium surge (Figures 8B and 8C, Video 7). Considering that Annexin V-binding to PS and accumulation on dendrite surface are likely slower than calcium activation of GCaMP6s, our data support the idea that PS-mediated phagocytosis causes dendrite membrane rupture and the final calcium surge.

Because the unique pattern of calcium flashing is absent in uninjured dendrites and *Nmnat* KO neurons, we suspected that it may play an active role in promoting degeneration of injured dendrites. If so, factors that can block dendrite degeneration may also alter the calcium flashing. Indeed, Wld^S^ OE dramatically reduced calcium fluctuations in injured dendrites and eliminated the quiescent and surge phases for the entire duration of our time-lapse imaging (13 hrs) (Figures 8E, 8J, S3B, and Video 8). In addition, using time-lapse imaging at a higher temporal resolution (2 s/frame), we found that wildtype injured dendrites displayed calcium flashes at a frequency of 0.4-3/min (Figures 8G and 8K, and Video 9) while injured dendrites of Wld^S^ OE neurons either maintained a much milder calcium fluctuation or exhibited irregular and infrequent calcium flashes (Figures 8H and 8K–8L; Videos 10 and 11). In contrast, the majority of severed dendrites of *axed* KO neurons (6/9) maintained constant and high calcium levels before the quiescence and surge phases (Figures 8F, 8J, S3C, and Video 12), which was further confirmed by time-lapse imaging at the high temporal resolution (Figures 8I, 8K, 8L, S3C, and Video 13). Overall, injured dendrites of Wld^S^ OE and *axed* KO neurons showed much smaller average amplitudes of calcium fluctuations compared to the wildtype control (Figure 8L). Because Wld^S^ OE and *axed* KO did not alter calcium dynamics in uninjured dendrites (Figures S3E and S3F), these data suggest that these manipulations specifically modify calcium flashing patterns in injured dendrites, consistent with the idea that calcium flashes may promote degeneration of injured dendrites.

## DISCUSSION

In this study, we investigate the mechanism of dendrite degenerations caused by genetic NAD^+^disruptions and injury in the *Drosophila* PNS. Although neurite phagocytosis has been observed after neuronal injury both *in vivo* and *in vitro* (Han et al., 2014; MacDonald et al., 2006; Nomura-Komoike et al., 2020; Sapar et al., 2018), WD is generally thought as a result of neurite self-destruction triggered by NAD^+^ depletion (Babetto et al., 2013; Gerdts et al., 2015; Gerdts et al., 2016; Neukomm et al., 2017). However, our results using the engulfment-deficient *drpr* mutant strongly suggest that PS-mediated phagocytosis is the main driving force for NAD^+^reduction-related dendrite degeneration *in vivo*. It is solely responsible for dendrite degeneration of *Nmnat* KO neurons and greatly accelerates the degeneration of Sarm^GOF^ OE neurons. In injury, phagocytosis is responsible for at least half of the dendrite fragmentation by 10 hrs AI and may contribute more to dendrite breakdown than self-destruction at earlier stages.

Supporting this idea, ectopic PS exposure on injured dendrites is sufficient to revert the blockage of dendrite fragmentation by Wld^S^ OE or *Sarm* KO, even though ectopically induced PS exposure is much lower than natural PS exposure on injured dendrites (Sapar et al., 2018). Our time-lapse analyses of PS exposure, the final calcium surge, and dendrite rupture also support that PS-mediated phagocytosis breaks down injury dendrites at the time of dendrite fragmentation. Therefore, at least in the context of dendrite injury, phagocytosis is the major factor driving degeneration while self-destruction acts as a secondary mechanism to ensure complete fragmentation.

NAD^+^ reduction is known to be essential for neuronal PS exposure and neurite self-destruction during WD (Gerdts et al., 2015; Sapar et al., 2018; Shacham-Silverberg et al., 2018). How does the same signaling input coordinate the two different events? Our results show that NAD^+^ disruption controls PS exposure and neurite self-destruction in separate steps of WD. In the existing model, Sarm activation is believed to cause catastrophic NAD^+^ depletion that is sufficient to initiate neurite self-destruction (Gerdts et al., 2015; Sasaki et al., 2016). However, we found that downstream of Sarm activation and before the initiation of self-destruction, neurites first expose PS to engage in phagocytosis-mediated non-autonomous degeneration. Therefore, in our revised model, dendrites respond to at least three distinct, increasingly severe levels of NAD^+^ reduction by eliciting different molecular events (Figure 9). Between the NAD^+^ level required for Sarm activation (SA level) and the level that initiates self-destruction (SD level), Sarm activity lowers NAD^+^ to a level that causes neurons to expose PS on their surface (which we call the PSE level). This PS exposure is sufficient to cause phagocytosis-mediated dendrite degeneration, which can be completely prevented by blocking engulfment activity of phagocytes. However, below the SD level, neurites spontaneously fragment even in the absence of phagocytosis.

**Figure 9.**
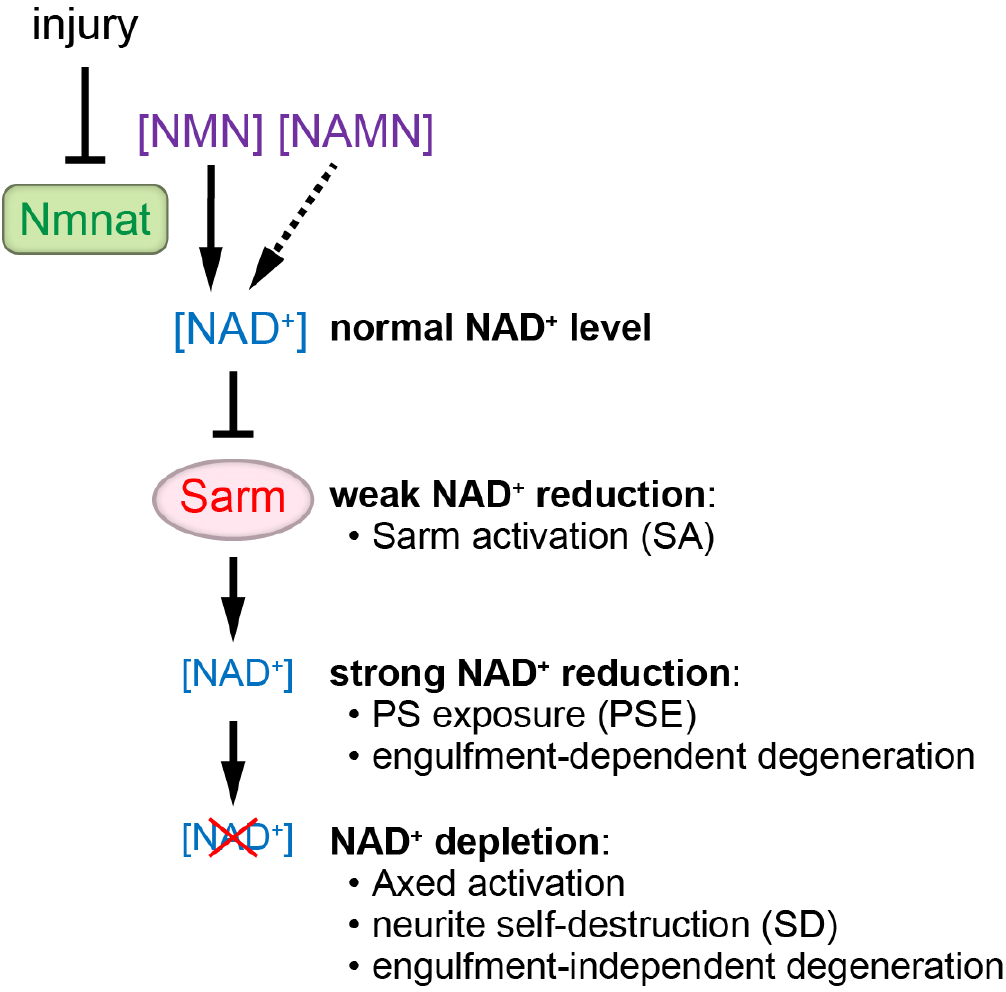
Three levels of NAD^+^ reduction elicit distinct responses in dendrites. See Discussion for details.

Our results also suggest a direct correlation between the kinetics of NAD^+^ reduction and the severity of neurite degeneration. *Nmnat* KO is expected to cause slow NAD^+^ reduction, due to gene perdurance and the time required for natural NAD^+^ turnover, and correspondingly causes engulfment-dependent dendrite degeneration only in late 3^rd^ instar larvae. In contrast, Sarm^GOF^ OE should lead to a more rapid NAD^+^ depletion and in fact causes engulfment-dependent dendrite degeneration as early as the 1^st^ instar and dendrite self-destruction by the 3^rd^ instar. Injury is known to cause even more rapid NAD^+^ reduction in axons (Wang et al., 2005) and is correlated with the fastest dendrite degeneration – initiation at around 4 hrs AI and completion usually by 10 hrs AI.

How does NAD^+^ reduction cause PS exposure? A direct consequence of NAD^+^ loss is the decline of neurite ATP levels due to the requirement of NAD^+^ in glycolysis and oxidative phosphorylation (Shacham-Silverberg et al., 2018; Wang et al., 2005). Consistent with ATP reduction playing a role in inducing PS exposure, suppressing mitochondria ATP synthesis in DRG culture caused gradual axonal PS exposure (Shacham-Silverberg et al., 2018). However, how ATP reduction may induce PS exposure remain elusive. Although the maintenance of membrane PS asymmetry by flippases requires ATP, flippase KO in C4da neurons causes a much milder PS exposure than injury (Sapar et al., 2018), suggesting that mechanisms other than flippase inhibition must be contributing to the rapid PS exposure seen after injury. Identifying the PS transporters responsible for PS exposure on injured neurites will be a key step for revealing the mechanisms of NAD^+^ regulation of PS exposure.

Genetic analyses in *Drosophila* identified Axed as a key switch of WD, whose activity is absolutely required for axon degeneration caused by injury and genetic depletion of NAD^+^(Neukomm et al., 2017). How does Axed regulate neurite degeneration? Our data suggest that Axed is not required for PS-mediated phagocytosis but contributes to self-destruction of injured dendrites, placing its activation below the SD level of NAD^+^ (Figure 9). Surprisingly, Axed seems to play a minor role in dendrite degeneration, as its LOF only slowed down but did not block self-destruction, indicating the existence of other factors that promote self-destruction of injured dendrites. Importantly, these results suggest that dendrites and axons may rely on different mechanisms of self-destruction.

By exploring the different mechanisms employed in *Nmnat* KO- and injury-induced dendrite degenerations, we discovered dynamic calcium activities only present in injured dendrites, including an unreported calcium flashing pattern prior to any obvious degenerative event and a final calcium surge that coincides with dendrite fragmentation. Calcium surge at the time of neurite fragmentation is a shared feature between injured axons of zebrafish (Vargas et al., 2015) and injured dendrites of *Drosophila* da neurons. Although calcium influx is required for WD (George et al., 1995) and may activate calcium-dependent lipid scramblases (Suzuki et al., 2010), our time-lapse analyses suggest that the final calcium surge is more likely a result of phagocytosis-induced membrane rupture rather than the cause of fragmentation. In comparison, the calcium flashing soon after the injury is unique to dendrites and may play an active role in dendrite degeneration in ways similar to the compartmentalized calcium flashing that occurs during developmental pruning of C4da neurons (Kanamori et al., 2013). Consistent with this possibility, the calcium flashing is suppressed by Wld^S^ OE and *axed* KO. Interestingly, these two manipulations block the calcium flashing in opposite ways, with Wld^S^ OE dampening the calcium level and *axed* KO keeping the calcium level high. This distinction may be a useful clue for understanding the regulation of PS exposure and self-destruction. For example, it is possible that elevated calcium levels prepare dendrites for PS exposure and drastic changes of calcium levels promote dendrite self-destruction.

Lastly, as impairment of NAD^+^ metabolism is a general feature of neurodegenerative disorders including Leber congenital amaurosis (LCA), Alzheimer’s disease, Parkinson’s disease, and retinal degenerations (Ali et al., 2013; Canto et al., 2015; Fang et al., 2017; Lin et al., 2016; Verdin, 2015), phagocytosis may play important roles in the pathogenesis of these diseases through dysregulated neuronal PS exposure.

## METHODS

### Fly strains

The details of fly strains used in this study are listed in the key reagent table. For labeling of C4da neurons, we used *ppk-CD4-tdTom, ppk-MApHS*, and *ppk-Gal4 UAS-CD4-tdTom*. For labeling PS exposure on dendrites, we used *dcg-Gal4 UAS-GFP-LactC1C2, R16A03-LexA LexAop-GFP-LactC1C2*, and *dcg-Gal4 UAS-AnnexinV-mCard*. To label Or22a axons, we used *Or22a-Gal4 UAS-mCD8-GFP*. To visualize dendrite rupture, we used *dcg-Gal4 UAS-sGFP(11)x7* together with *ppk-LexA LexAop-myr-tdTom-GFP(1-10)*. To visualize calcium activities in C4da dendrites, we used *ppk-LexA LexAop-myr-GCaMP6s*.

### Molecular cloning and transgenic flies

#### ey-Cas9

Three tandem copies of a 211bp *ey* enhancer (corresponding to nucleotides 2577-2787 in GenBank accession number AJ131630) was inserted into XhoI/SalI sites of pENTR11 (Thermo Fisher Scientific). The resulting entry vector was combined with a Cas9 destination vector which is similar to pDEST-APIC-Cas9 (Addgene 121657) but does not contain Inr, MTE, and DPE in the Hsp70 core promoter (Poe et al., 2019) to generate the pAPIC-ey-Cas9 expression vector through a Gateway LR reaction.

#### LexAop-myr-tdTom-GFP(1-10)

pAPLO-CD4-tdTom (Poe et al., 2017) was digested by BglII/AscI, blunted, and religated to remove an XbaI site before 13xLexAop2. The resulting construct is called pAPLOm-CD4-tdTom. An myr-GFP fragment was isolated from pJFRC176-10XUAS-rox-dSTOP-rox-myr-GFP (Addgene 32147) by XhoI/XbaI digestion and ligated to XhoI/XbaI sites of pAPLOm-CD4-tdTom to make pAPLO-myr-GFP. pAPLO-myr-GFP was then digested by BamHI/XbaI and subsequently assembled with a tdTom PCR fragment and a GFP(1-10) gBlock fragment (synthesized by IDT) through NEBuilder DNA Assembly to generate pAPLO-myr-tdTom-GFP1-10.

#### UAS-sGFP(11)x7

A DNA fragment containing GFP(11)x7 was PCR amplified from a gBlock DNA fragment and cloned into Nhe/XbaI sites of pIHEU-sfGFP-LactC1C2 using NEBuilder DNA Assembly. The resulting construct pIHEU-sGFP11×7 inherits the signal peptide sequence from pIHEU-sfGFP-LactC1C2 but removes the sfGFP-LactC1C2 coding sequence.

#### LexAop-WldS

The WldS coding sequence was PCR amplified from UAS-WldS genomic DNA and cloned into XhoI/XbaI sites of pAPLOm-CD4-tdTom via restriction cloning to make pAPLO-WldS.

#### UAS-ATP8A

A fragment encoding a FLAG tag and the ER exit signal of Kir2.1 (Han et al., 2011) was cloned into the EcoRI and XbaI sites of pACU (Han et al., 2011) by ligating with annealed oligos. The ATP8A coding region that is shared by all isoforms was amplified by PCR from cDNA clone GH28327 (AA26 to AA1091, BDGP) and inserted into an NheI site before the FLAG tag.

#### gRNA expression vectors

For *Nmnat, Sarm*, and *axed*, gRNAs were cloned into pAC-U63-tgRNA-Rev as described (Poe et al., 2019). The first tRNA spacer in each final construct is tRNA^Gly^ but the remaining tRNA spacers are tRNA^Gln^ (Koreman et al., 2021). *gRNA-peb* was constructed in pAC-U63-QtgRNA2.1-BR. The gRNA target sequences are listed in the table below.

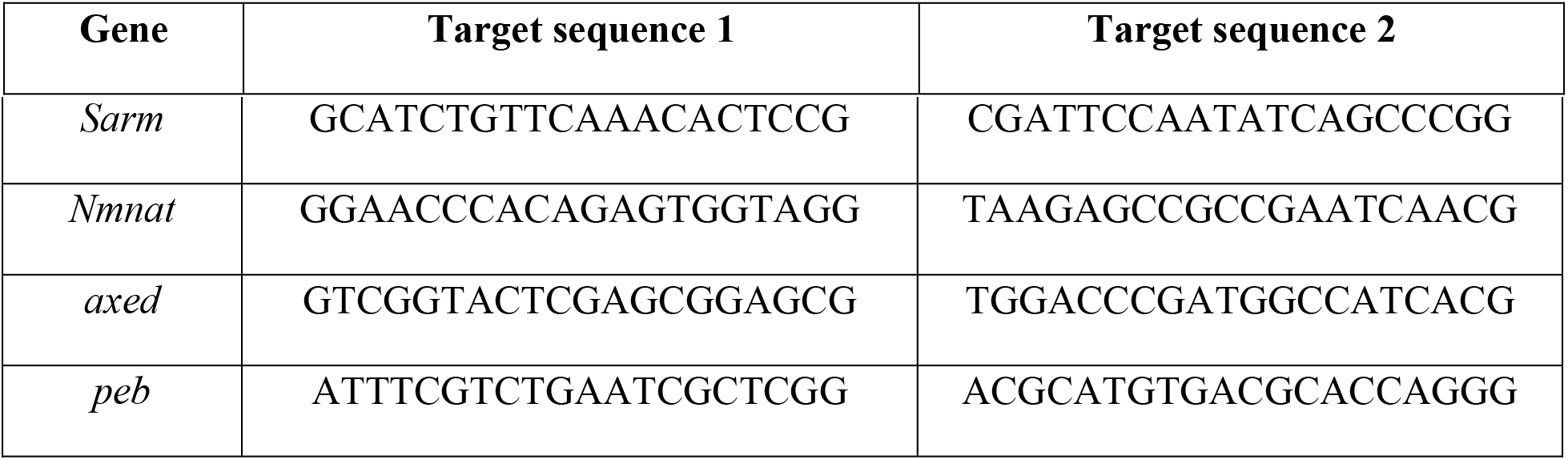

The above constructs were injected by Rainbow Transgenic Flies to transform flies through φC31 integrase-mediated integration into attP docker sites. The sequences of constructs will be provided upon request.

### CRISPR-TRiM

The efficiency of transgenic gRNA lines was validated by the Cas9-LEThAL assay (Poe et al., 2019). Homozygous males of each gRNA line were crossed to *Act-Cas9 w lig4* homozygous females. *gRNA-Nmnat* crosses caused lethality of all progeny at the 2^nd^ instar larval stage; *gRNA-Sarm* crosses yielded viable female progeny and male lethality between 3^rd^ instar larvae to prepupae; *gRNA-axed* crosses caused lethality at the embryonic stage for all progeny; *gRNA-peb* crosses caused lethality from embryonic stage to the 2^nd^ instar for all progeny. These results suggest that all gRNAs are efficient.

C4da-specific gene knockout was carried out using *ppk-Cas9* (Poe et al., 2019). Tissue-specific knockout in da neuron precursor cells were carried out with *SOP-Cas9* (Poe et al., 2019). Gene knockout in the precursor cells of Or22a ORNs was carried out using *ey-Cas9* (this study).

### Live imaging

Animals were reared at 25°C in density-controlled vials (60-100 embryos/vial) on standard yeast-glucose medium (doi:10.1101/pdb.rec10907). Larvae at 125 hours AEL (wandering stage) or stages specified were mounted in 100% glycerol under coverslips with vacuum grease spacer and imaged using a Leica SP8 microscope equipped with a 40X NA1.30 oil objective. Larvae were lightly anesthetized with isoflurane before mounting. For consistency, we imaged dorsal ddaC neurons from A1-A3 segments (2-3 neurons per animal) on one side of the larvae. Unless stated otherwise, confocal images shown in all figures are maximum intensity projections of z stacks encompassing the epidermal layer and the sensory neurons beneath, which are typically 8–10 μm for 3^rd^ instar larvae.

#### Injury assay

Injury assay at the larval stage was done as described previously (Sapar et al., 2018). Briefly, larvae at 90 hr AEL were lightly anesthetized with isoflurane, mounted in a small amount of halocarbon oil under coverslips with grease spacers. The laser ablation was performed on a Zeiss LSM880 Confocal/Multiphoton Upright Microscope, using a 790 nm two-photon laser at primary dendrites of ddaC neurons in A1 and A3 segments. Animals were recovered on grape juice agar plates following lesion for appropriate times before imaging.

ORN axon injury assay was performed on 7-day-old male flies by removing the outer segments of both antennae as described in (MacDonald et al., 2006). The injured males were recovered for 7 days by transferring to fresh yeast-glucose medium every day. To examine Or22a axon degeneration, brains were dissected in PBST (0.2% Triton-X in PBS), fixed in 4% formaldehyde in PBS for 20 min, and then rinsed with PBST three times, 20 minutes each. Then the brains were mounted in SlowFade^®^ Diamond Antifade Mountant (Thermo Fisher Scientific) and imaged using a Leica SP8 microscope with a 40x NA1.3 oil objective.

#### Long-term time-lapse imaging

Long-term time-lapse imaging at the larval stage was done as described previously (Ji and Han, 2020; Sapar et al., 2018). Briefly, a layer of double-sided tape was placed on the coverslip to define the position of PDMS blocks. A small amount of UV glue was added to the groove of PDMS and to the coverslip. Anesthetized larvae were placed on top of the UV glue on the coverslip and then covered by PDMS blocks with the groove side contacting the larva. Glue was then cured by 365nm UV light. The coverslip with attached PDMS and larvae was mounted on an aluminum slide chamber that contained a piece of moisturized Kimwipes (Kimtech Science) paper. Time-lapse imaging was performed on a Leica SP8 confocal equipped with a 40x NA1.3 oil objective and a resonant scanner at digital zoom 0.75 and a 3-min or 2-sec interval. For imaging after ablation, larvae were pre-mounted in the imaging chamber and subjected to laser injury. The larvae were then imaged 0.5-1 hours after ablation. For calcium imaging before and immediately after ablation, images were captured on a Zeiss LSM880 Confocal/Multiphoton Upright Microscope on which the ablation was performed.

### Image analysis and quantification

Image processing and analyses were done in Fiji/ImageJ. Methods for tracing and measuring C4da neuron dendrite length have been previously described (Poe et al., 2017). Briefly, the images were segmented by Auto Local Threshold and reduced to single pixel skeletons before measurement of skeleton length by pixel distance. The dendrite debris measurement has been described previously (Sapar et al., 2018). Briefly, a dendrite mask was first generated from projected images by Auto Local Threshold in order to create a region of interest (ROI) by dilation to map areas within one-epidermal-cell diameter (40 μm) from dendrites. Dendrite debris within the ROI was converted to binary masks based on fixed thresholds. Different thresholds were used for *ppk-C4-tdTom* and *ppk-Gal4 UAS-CD-tdTom* as they have different brightness. The dendrite pixel area (ADen), debris pixel area (ADeb), and ROI area (AROI) were measured and dendrite coverage ratio was calculated based on the following formula: 100·Adeb·AROI /(AROI-ADen)·ADen. For measuring Lact-GFP, two regions at empty epidermal regions were measured as background levels. TdTom signals on dendrites were used to generate dendrite masks for measurement of GFP within the masks. For kymographs, we used a custom macro based on the Straighten function to extract a strip of pixels centered at the selected dendrite branch. The maximum intensity pixel in the strip at each distance was used to generate a single-pixel line for each time frame. The final kymographs were displayed using the Fire lookup table (LUT).

### Statistical Analysis

R was used to conduct statistical analyses and generate graphs. (*p < 0.05, **p < 0.01, and ***p < 0.001). Statistical significance was set at p < 0.05. Data acquisition and quantification were performed non-blinded. Acquisition was performed in ImageJ (batch processing for debris coverage ratio and fragmentation ratio, manually by hand for GFP-Lact binding) and Microsoft Excel. Statistical analyses were performed using R. We used the following R packages: car, stats, multcomp for statistical analysis and ggplot2 for generating graphs. Some graphs were made in Excel using its native plotting functions. For the statistical analysis we ran the following tests, ANOVA (followed by Tukey’s HSD) when dependent variable was normally distributed and there was approximately equal variance across groups. When dependent variable was not normally distributed and variance was not equal across groups, we used Kruskal-Wallis (followed by Dunn’s test, p-values adjusted with Benjamini-Hochberg method) to test the null hypothesis that assumes that the samples (groups) are from identical populations. To check whether the data fit a normal distribution, we generated qqPlots to analyze whether the residuals of the linear regression model is normally distributed. We used the Levene’s test to check for equal variance within groups. The state of neuronal degeneration or fragmentation was compared using the Freeman-Halton extension of Fisher’s exact test.

#### Replication

For all larval and adult imaging experiments, at least 3 biological replications were performed for each genotype and/or condition.

## AUTHOR CONTRIBUTIONS

Conceptualization, CH, HJ, MLS, AS; Methodology, CH, HJ, MLS, AS, BW; Investigation, HJ, MLS, AS; Formal Analysis, HJ, MLS, AS; Resources, CH, MLS, HJ, BW, AS; Writing – Original Draft, CH, MLS, HJ, AS; Writing – Review and Editing, CH, MLS, HJ, AS; Funding Acquisition, CH.

## ACKNOWLEDGMENTS

We thank Marc Freeman, Yang Xiang, Heather Broihier, and Bloomington Stock Center for fly stocks; Addgene for plasmids; Cornell BRC Imaging facility for access to microscopes (funded by NIH grant S10OD018516); Cornell CSCU for advice on statistics; Mike Goldberg and Fenghua Hu for critical reading and suggestions on the manuscript. This work was supported by a Cornell start-up fund and NIH grants (R01NS099125 and R21OD023824) awarded to C.H.

---

**Figure S1:**
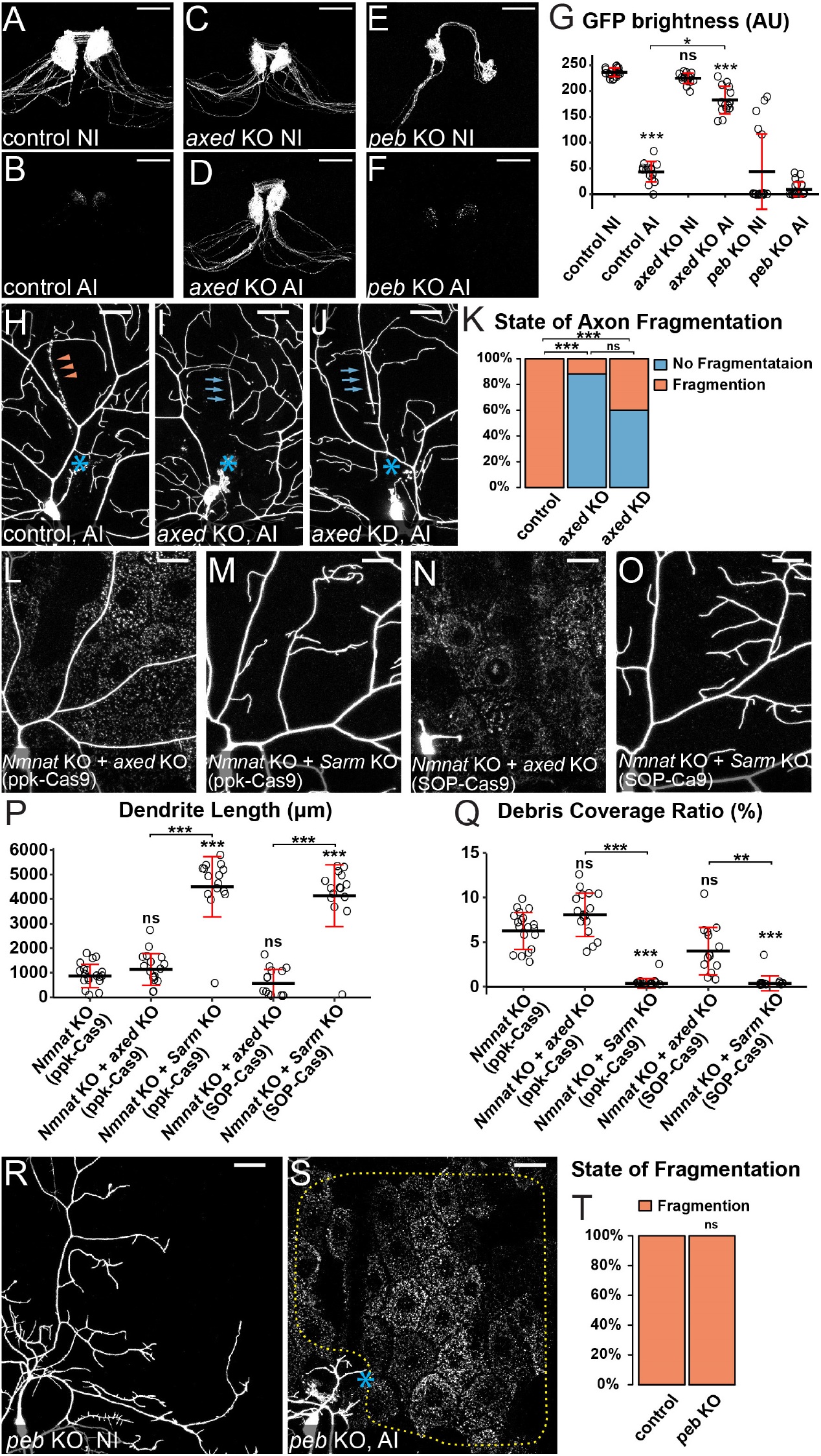
Axed is not involved in PS-mediated phagocytosis. (A-F) Axons of control NI (A), control 7 days AI (B), *axed* KO NI (C), *axed* KO 7 days AI (D), *peb* KO NI (E), and *peb* KO 7 days AI (F) Or22a neurons in 14-day-old adult brains. Or22a neurons were labeled by *Or22a>CD8-GFP* and KO was induced by *ey-Cas9*. Scale bars, 25 μm. (G) Quantification of GFP brightness of Or22a glomeruli. n = number of brains: control NI 14 days (n = 19); control 7 days AI 14 days (n = 15); *axed* KO NI 14 days (n = 16); *axed* KO 7 days AI 14 days (n = 14); *peb* KO NI 14 days (n = 18); *peb* KO 7 days AI 14 days (n = 20). Kruskal-Wallis (One-way ANOVA on ranks) and Dunn’s test, p-values adjusted with the Benjamini-Hochberg method. *p≤0.05, ***p<0.001, n.s., not significant. The significance level above each genotype is for comparison with the control. Statistical comparisons for *peb* KO glomeruli are not shown due to the variation within the group of *peb* KO NI 14 days. Black bar, mean; red bar, SD. (H-J) Injured axons of wildtype 24 hrs AI (H), *axed* KO AI 24 hrs AI (I), and *axed* KD AI 24 hrs AI (J). Neurons were labeled by *ppk-MApHS*. Neuronal-specific KO was induced by *SOP-Cas9*, and neuronalspecific KD was driven by *21-7 Gal4*. Scale bars, 25 μm. (K) Quantification of axon fragmentation showing percentages of neurons undergoing no fragmentation, and complete fragmentation of injured axons at 24 hrs AI. n = number of neurons: wildtype 24 hrs AI (n = 20, 12 animals); *axed* KO 24 hrs AI (n = 17, 9 animals); *axed* KD 24hrs AI (n = 20, 8 animals). Freeman-Halton extension of Fisher’s exact test. For all quantifications, ***p≤0.001; n.s., not significant. (L-O) Partial dendritic fields of *Nmnat* KO + *axed* KO (L and N) and *Nmnat* KO + *Sarm* KO (M and O) ddaC C4da neurons. Neurons were labeled by *ppk>CD4-tdTom* (L-O). The C4da-specific KO was induced by *ppk-Cas9* (L-M) or by *SOP-Cas9* (N-O). Scale bars, 25 μm. (P) Quantification of dendrite length. n = number of neurons: *Nmnat* KO (*ppk-Cas9*) (n = 19, 10 animals); *Nmnat* KO + *axed* KO (*ppk-Cas9*) (n = 17, 9 animals); *Nmnat* KO +*Sarm* KO (*ppk-Cas9*) (n = 16, 8 animals); *Nmnat* KO + *axed* KO (*SOP-Cas9*) (n = 14, 7 animals); *Nmnat* KO + *Sarm* KO (*SOP-Cas9*) (n = 15, 8 animals). One-way ANOVA and Tukey’s test. (Q) Quantification of debris coverage ratio, which is the percentage of debris area normalized by dendrite area ratio. Number of neurons: same as in (P). Kruskal-Wallis (One-way ANOVA on ranks) and Dunn’s test, p-values adjusted with the Benjamini-Hochberg method. For all quantifications, **p≤0.01; ***p≤0.001; n.s., not significant. The significance level above each genotype is for comparison with the control. Black bar, mean; red bar, SD. (R-S) Partial dendritic fields of *peb* KO NI (R) and *peb* KO 24 hrs AI (S) ddaC neurons. Yellow dots outline regions covered by injured dendrites in (S). Neurons were labeled by *ppk>CD4-tdTom* and neuronal-specific KO was induced by *SOP-Cas9*. Scale bars, 25 μm. (T) Quantification of dendrite fragmentation showing percentages of neurons undergoing complete fragmentation of injured dendrites at 24 hrs AI. n = number of neurons: wildtype 24 hrs AI (n = 21, 12 animals); *peb* KO 24 hrs AI (n = 12, 7 animals). Freeman-Halton extension of Fisher’s exact test. For all quantifications, n.s., not significant.

**Figure S2:**
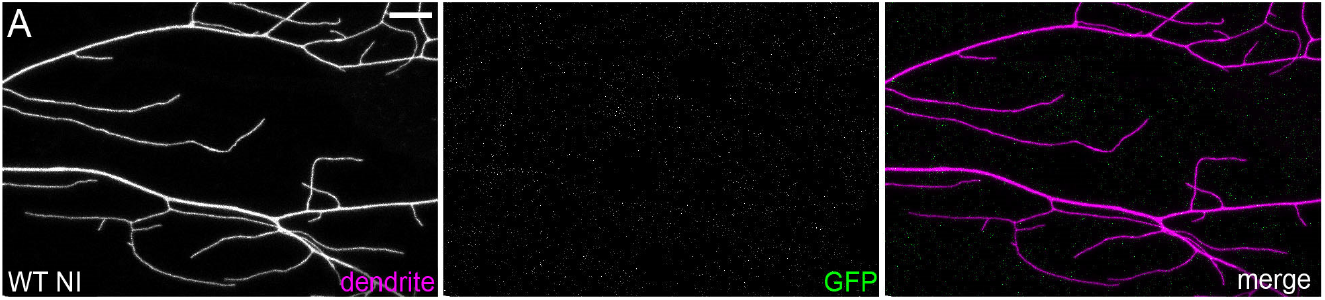
Membrane of uninjured dendrites does not rupture. (A) Uninjured wildtype dendrites lacking reconstituted GFP in the membrane rupture assay. Scale bar, 25 μm.

**Figure S3:**
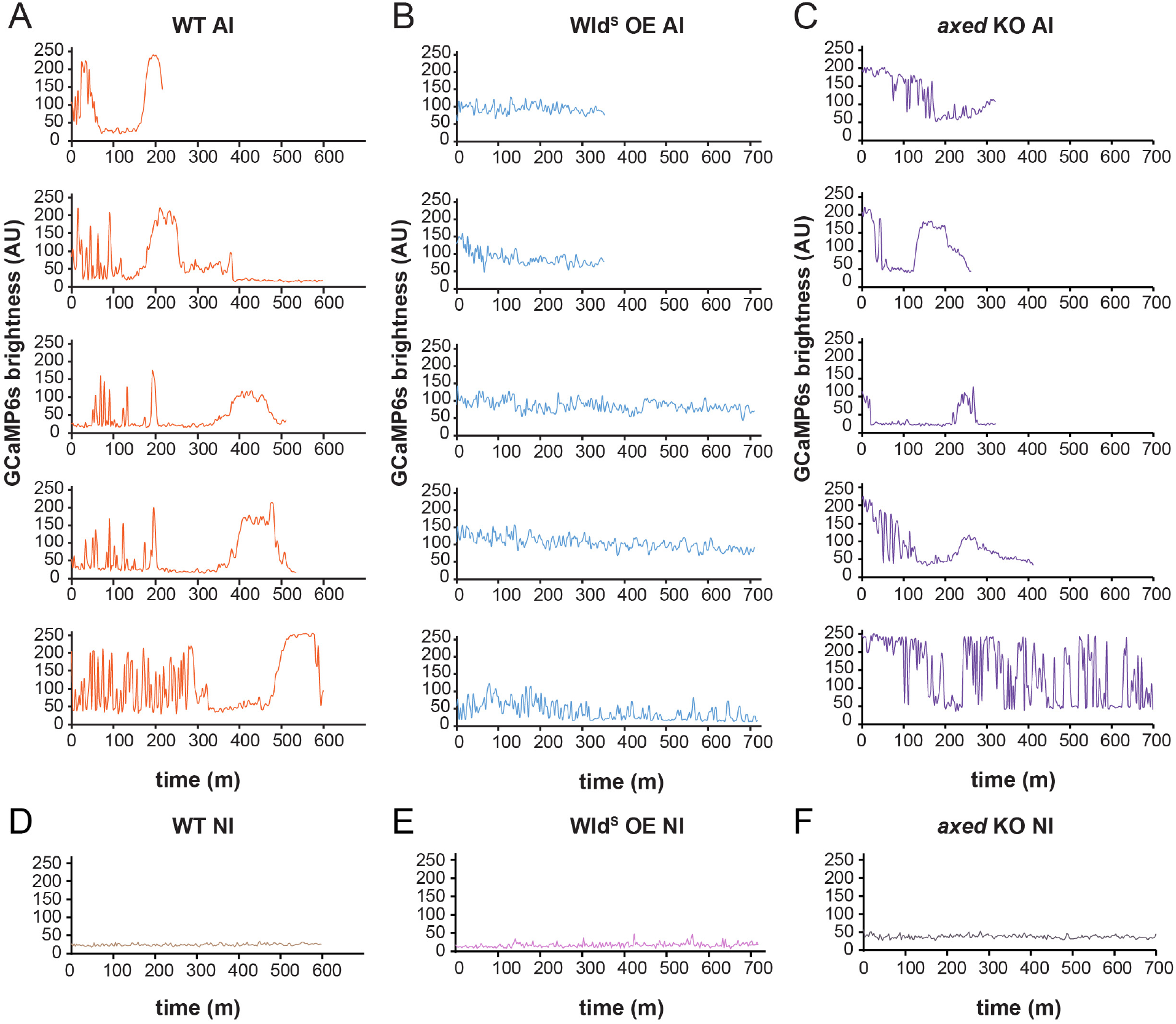
Wld^S^ overexpression and loss of *axed* changed calcium flashes in injured dendrites. (A-C) Representative plots of GCaMP6s brightness in injured dendrites of wildtype (A), Wld^S^-overexpressing (B), and *axed* KO (C) neurons over time with 3-min interval. (D-F) Representative plots of GCaMP6s brightness in uninjured dendrites of wildtype (A), Wld^S^-overexpressing (B), and *axed* KO (C) neurons over time with 3-min interval.

## VIDEO LEGEND

**Video 1. GFP-Lact labeling of degenerating *Nmnat* KO dendrites, related to Figure 2.**

Time-lapse movie of GFP-Lact labeling on distal dendrites of a *Nmnat* KO C4da neuron before and during degeneration. Imaging started around 96 hours AEL. *Dcg-Gal4* drives GFP-Lact expression in both fat bodies (not shown) and hemocytes (mobile cells in the GFP-Lact channel). Timestamp is relative to the first frame, with a 3-min interval between each frame.

**Video 2. Dendrite dynamics of *Nmnat* KO neurons in *drpr* mutant larvae, related to Figure 2.**

Time-lapse movie of dendrites of a *Nmnat* KO neuron exhibiting dynamic extension and retraction behaviors in a *drpr* mutant larva. Imaging started around 120 hours AEL. Timestamp is relative to the first frame, with a 3-min interval between each frame.

**Video 3. Membrane rupture of injured dendrites, related to Figure 7.**

Time-lapse movie of laser-injured C4da dendrites from 1 to 11 hours AI. Reconstituted GFP is from extracellular GFP(11)x7 entering the dendrites and myr-tdTom-GFP(1-10) attached to the inner leaflet of dendritic membrane. Timestamp is relative to the first frame, with a 3-min interval between each frame.

**Video 4. Cytoplasmic calcium dynamics in uninjured wildtype dendrites, related to Figure 8.**

Time-lapse movie of an uninjured wildtype C4da neuron showing low baseline GCaMP6s signals and occasional local rises in dendrites. Timestamp is relative to the first frame, with a 3-min interval between each frame.

**Video 5. Cytoplasmic calcium dynamics in dendrites of *Nmnat* KO neurons, related to Figure 8.**

Time-lapse movie of a *Nmnat* KO C4da neuron showing calcium dynamics in uninjured dendrites. Timestamp is relative to the first frame, with a 3-min interval between each frame.

**Video 6. Calcium dynamics in injured wildtype dendrites, related to Figure 8.**

Time-lapse movie of laser-injured C4da dendrites from 1 to 11 hrs AI showing calcium dynamics in both severed dendrites and those attached to the soma. Timestamp is relative to the first frame, with a 3-min interval between each frame.

**Video 7. AV-mCard labeling and calcium dynamics of injured dendrites, related to Figure 8.**

Time-lapse movie of laser-injured C4da dendrites from 1 to 6 hrs AI showing labeling of injured dendrites by the PS sensor AV-mCard and GCaMP6s signals. Timestamp is relative to the first frame, with a 3-min interval between each frame.

**Video 8. Calcium dynamics in injured Wld^S^ OE dendrites, related to Figure 8.**

Time-lapse movie of laser-injured Wld^S^ OE C4da dendrites from 1 to 13 hours AI showing GCaMP6s signals in severed dendrites. Timestamp is relative to the first frame, with a 3-min interval between each frame.

**Video 9. Calcium dynamics in injured wildtype dendrites with a higher temporal resolution, related to Figure 8.**

High temporal-resolution time-lapse movie of laser-injured wildtype C4da dendrites around 1 hr AI showing calcium dynamics. Timestamp is relative to the first frame, with a 2-sec interval between each frame.

**Video 10. Calcium dynamics in injured Wld^S^ OE dendrites at a higher temporal resolution, related to Figure 8.**

High temporal-resolution time-lapse movie of laser-injured Wld^S^ OE C4da dendrites around 2 hrs AI showing calcium dynamics. Timestamp is relative to the first frame, with a 2-sec interval between each frame.

**Video 11. Irregular and infrequent cytoplasmic calcium flashes in injured Wld^S^ OE dendrites at a higher temporal resolution, related to Figure 8.**

High temporal-resolution time-lapse movie of laser-injured Wld^S^ OE C4da dendrites around 2 hrs AI showing irregular and infrequent GCaMP6s signals in severed dendrites. Timestamp is relative to the first frame, with a 2-sec interval between each frame.

**Video 12. Calcium dynamics in injured *axed* KO dendrites, related to Figure 8.**

Time-lapse movie of laser-injured *axed* KO C4da dendrites from 0.5 to 5.5 hours AI showing GCaMP6s signals in severed dendrites. Timestamp is relative to the first frame, with a 3-min interval between each frame.

**Video 13. Calcium dynamics in injured *axed* KO dendrites at a higher temporal resolution, related to Figure 8.**

High temporal-resolution time-lapse movie of laser-injured *axed* KO C4da dendrites around 10 min AI showing calcium dynamics. Timestamp is relative to the first frame, with a 2-sec interval between each frame.

## REFERENCE

Akbergenova, Y., Cunningham, K.L., Zhang, Y.V., Weiss, S., and Littleton, J.T. (2018). Characterization of developmental and molecular factors underlying release heterogeneity at Drosophila synapses. Elife 7.

Ali, Y.O., Li-Kroeger, D., Bellen, H.J., Zhai, R.G., and Lu, H.C. (2013). NMNATs, evolutionarily conserved neuronal maintenance factors. Trends Neurosci 36, 632–640.

Avery, M.A., Sheehan, A.E., Kerr, K.S., Wang, J., and Freeman, M.R. (2009). Wld S requires Nmnat1 enzymatic activity and N16-VCP interactions to suppress Wallerian degeneration. J Cell Biol 184, 501–513.

Babetto, E., Beirowski, B., Russler, E.V., Milbrandt, J., and DiAntonio, A. (2013). The Phr1 ubiquitin ligase promotes injury-induced axon self-destruction. Cell Rep 3, 1422–1429.

Bratkowski, M., Xie, T., Thayer, D.A., Lad, S., Mathur, P., Yang, Y.S., Danko, G., Burdett, T.C., Danao, J., Cantor, A., et al. (2020). Structural and Mechanistic Regulation of the Pro-degenerative NAD Hydrolase SARM1. Cell Rep 32, 107999.

Canto, C., Menzies, K.J., and Auwerx, J. (2015). NAD(+) Metabolism and the Control of Energy Homeostasis: A Balancing Act between Mitochondria and the Nucleus. Cell Metab 22, 31–53.

Coleman, M.P., and Freeman, M.R. (2010). Wallerian degeneration, wld(s), and nmnat. Annu Rev Neurosci 33, 245–267.

Davies, A.J., Kim, H.W., Gonzalez-Cano, R., Choi, J., Back, S.K., Roh, S.E., Johnson, E., Gabriac, M., Kim, M.S., Lee, J., et al. (2019). Natural Killer Cells Degenerate Intact Sensory Afferents following Nerve Injury. Cell 176, 716–728 e718.

Di Stefano, M., Nascimento-Ferreira, I., Orsomando, G., Mori, V., Gilley, J., Brown, R., Janeckova, L., Vargas, M.E., Worrell, L.A., Loreto, A., et al. (2015). A rise in NAD precursor nicotinamide mononucleotide (NMN) after injury promotes axon degeneration. Cell Death Differ 22, 731–742.

Fang, E.F., Lautrup, S., Hou, Y., Demarest, T.G., Croteau, D.L., Mattson, M.P., and Bohr, V.A. (2017). NAD(+) in Aging: Molecular Mechanisms and Translational Implications. Trends Mol Med 23, 899–916.

Farley, J.E., Burdett, T.C., Barria, R., Neukomm, L.J., Kenna, K.P., Landers, J.E., and Freeman, M.R. (2018). Transcription factor Pebbled/RREB1 regulates injury-induced axon degeneration. Proc Natl Acad Sci U S A 115, 1358–1363.

Figley, M.D., Gu, W., Nanson, J.D., Shi, Y., Sasaki, Y., Cunnea, K., Malde, A.K., Jia, X., Luo, Z., Saikot, F.K., et al. (2021). SARM1 is a metabolic sensor activated by an increased NMN/NAD(+) ratio to trigger axon degeneration. Neuron 109, 1118–1136 e1111.

Fourgeaud, L., Traves, P.G., Tufail, Y., Leal-Bailey, H., Lew, E.D., Burrola, P.G., Callaway, P., Zagorska, A., Rothlin, C.V., Nimmerjahn, A., et al. (2016). TAM receptors regulate multiple features of microglial physiology. Nature 532, 240–244.

Freeman, M.R., Delrow, J., Kim, J., Johnson, E., and Doe, C.Q. (2003). Unwrapping glial biology: Gcm target genes regulating glial development, diversification, and function. Neuron 38, 567–580.

Galloway, D.A., Phillips, A.E.M., Owen, D.R.J., and Moore, C.S. (2019). Phagocytosis in the Brain: Homeostasis and Disease. Front Immunol 10, 790.

George, E., Glass, J., and Griffin, J. (1995). Axotomy-induced axonal degeneration is mediated by calcium influx through ion-specific channels. The Journal of Neuroscience 15, 6445–6452.

Gerdts, J., Brace, E.J., Sasaki, Y., DiAntonio, A., and Milbrandt, J. (2015). SARM1 activation triggers axon degeneration locally via NAD(+) destruction. Science 348, 453–457.

Gerdts, J., Summers, D.W., Milbrandt, J., and DiAntonio, A. (2016). Axon Self-Destruction: New Links among SARM1, MAPKs, and NAD+ Metabolism. Neuron 89, 449–460.

Han, C., Jan, L.Y., and Jan, Y.N. (2011). Enhancer-driven membrane markers for analysis of nonautonomous mechanisms reveal neuron-glia interactions in Drosophila. Proc Natl Acad Sci U S A 108, 9673–9678.

Han, C., Song, Y., Xiao, H., Wang, D., Franc, N.C., Jan, L.Y., and Jan, Y.N. (2014). Epidermal cells are the primary phagocytes in the fragmentation and clearance of degenerating dendrites in Drosophila. Neuron 81, 544–560.

Ji, H., and Han, C. (2020). LarvaSPA, A Method for Mounting Drosophila Larva for Long-Term Time-Lapse Imaging. J Vis Exp.

Jiang, Y., Liu, T., Lee, C.H., Chang, Q., Yang, J., and Zhang, Z. (2020). The NAD(+)-mediated self-inhibition mechanism of pro-neurodegenerative SARM1. Nature 588, 658–663.

Kanamori, T., Kanai, M.I., Dairyo, Y., Yasunaga, K., Morikawa, R.K., and Emoto, K. (2013). Compartmentalized calcium transients trigger dendrite pruning in Drosophila sensory neurons. Science 340, 1475–1478.

Koreman, G.T., Xu, Y., Hu, Q., Zhang, Z., Allen, S.E., Wolfner, M.F., Wang, B., and Han, C. (2021). Upgraded CRISPR/Cas9 tools for tissue-specific mutagenesis in Drosophila. Proc Natl Acad Sci U S A 118.

Leventis, P.A., and Grinstein, S. (2010). The distribution and function of phosphatidylserine in cellular membranes. Annu Rev Biophys 39, 407–427.

Lin, J.B., Kubota, S., Ban, N., Yoshida, M., Santeford, A., Sene, A., Nakamura, R., Zapata, N., Kubota, M., Tsubota, K., et al. (2016). NAMPT-Mediated NAD(+) Biosynthesis Is Essential for Vision In Mice. Cell Rep 17, 69–85.

Liu, H.W., Smith, C.B., Schmidt, M.S., Cambronne, X.A., Cohen, M.S., Migaud, M.E., Brenner, C., and Goodman, R.H. (2018). Pharmacological bypass of NAD(+) salvage pathway protects neurons from chemotherapy-induced degeneration. Proc Natl Acad Sci U S A.

MacDonald, J.M., Beach, M.G., Porpiglia, E., Sheehan, A.E., Watts, R.J., and Freeman, M.R. (2006). The Drosophila cell corpse engulfment receptor Draper mediates glial clearance of severed axons. Neuron 50, 869–881.

Mack, T.G., Reiner, M., Beirowski, B., Mi, W., Emanuelli, M., Wagner, D., Thomson, D., Gillingwater, T., Court, F., Conforti, L., et al. (2001). Wallerian degeneration of injured axons and synapses is delayed by a Ube4b/Nmnat chimeric gene. Nat Neurosci 4, 1199–1206.

Mazaheri, F., Breus, O., Durdu, S., Haas, P., Wittbrodt, J., Gilmour, D., and Peri, F. (2014). Distinct roles for BAI1 and TIM-4 in the engulfment of dying neurons by microglia. Nat Commun 5, 4046.

Miller, B.R., Press, C., Daniels, R.W., Sasaki, Y., Milbrandt, J., and DiAntonio, A. (2009). A dual leucine kinase-dependent axon self-destruction program promotes Wallerian degeneration. Nat Neurosci 12, 387–389.

Nandrot, E.F., Anand, M., Almeida, D., Atabai, K., Sheppard, D., and Finnemann, S.C. (2007). Essential role for MFG-E8 as ligand for alphavbeta5 integrin in diurnal retinal phagocytosis. Proc Natl Acad Sci U S A 104, 12005–12010.

Neukomm, L.J., Burdett, T.C., Seeds, A.M., Hampel, S., Coutinho-Budd, J.C., Farley, J.E., Wong, J., Karadeniz, Y.B., Osterloh, J.M., Sheehan, A.E., et al. (2017). Axon Death Pathways Converge on Axundead to Promote Functional and Structural Axon Disassembly. Neuron 95, 78–91 e75.

Nomura-Komoike, K., Saitoh, F., and Fujieda, H. (2020). Phosphatidylserine recognition and Rac1 activation are required for Muller glia proliferation, gliosis and phagocytosis after retinal injury. Sci Rep 10, 1488.

Osterloh, J.M., Yang, J., Rooney, T.M., Fox, A.N., Adalbert, R., Powell, E.H., Sheehan, A.E., Avery, M.A., Hackett, R., Logan, M.A., et al. (2012). dSarm/Sarm1 is required for activation of an injury-induced axon death pathway. Science 337, 481–484.

Poe, A.R., Tang, L., Wang, B., Li, Y., Sapar, M.L., and Han, C. (2017). Dendritic space-filling requires a neuronal type-specific extracellular permissive signal in Drosophila. Proc Natl Acad Sci U S A 114, E8062–E8071.

Poe, A.R., Wang, B., Sapar, M.L., Ji, H., Li, K., Onabajo, T., Fazliyeva, R., Gibbs, M., Qiu, Y., Hu, Y., et al. (2019). Robust CRISPR/Cas9-Mediated Tissue-Specific Mutagenesis Reveals Gene Redundancy and Perdurance in Drosophila. Genetics 211, 459–472.

Raiders, S., Black, E.C., Bae, A., MacFarlane, S., Klein, M., Shaham, S., and Singhvi, A. (2021). Glia actively sculpt sensory neurons by controlled phagocytosis to tune animal behavior. Elife 10.

Sapar, M.L., and Han, C. (2019). Die in pieces: How Drosophila sheds light on neurite degeneration and clearance. J Genet Genomics 46, 187–199.

Sapar, M.L., Ji, H., Wang, B., Poe, A.R., Dubey, K., Ren, X., Ni, J.Q., and Han, C. (2018). Phosphatidylserine Externalization Results from and Causes Neurite Degeneration in Drosophila. Cell Rep 24, 2273–2286.

Sasaki, Y., Nakagawa, T., Mao, X., DiAntonio, A., and Milbrandt, J. (2016). NMNAT1 inhibits axon degeneration via blockade of SARM1-mediated NAD(+) depletion. Elife 5.

Shacham-Silverberg, V., Sar Shalom, H., Goldner, R., Golan-Vaishenker, Y., Gurwicz, N., Gokhman, I., and Yaron, A. (2018). Phosphatidylserine is a marker for axonal debris engulfment but its exposure can be decoupled from degeneration. Cell Death Dis 9, 1116.

Shen, C., Vohra, M., Zhang, P., Mao, X., Figley, M.D., Zhu, J., Sasaki, Y., Wu, H., DiAntonio, A., and Milbrandt, J. (2021). Multiple domain interfaces mediate SARM1 autoinhibition. Proc Natl Acad Sci U S A 118.

Sievers, C., Platt, N., Perry, V.H., Coleman, M.P., and Conforti, L. (2003). Neurites undergoing Wallerian degeneration show an apoptotic-like process with Annexin V positive staining and loss of mitochondrial membrane potential. Neurosci Res 46, 161–169.

Suzuki, J., Umeda, M., Sims, P.J., and Nagata, S. (2010). Calcium-dependent phospholipid scrambling by TMEM16F. Nature 468, 834–838.

Tanaka, K., Fujimura-Kamada, K., and Yamamoto, T. (2011). Functions of phospholipid flippases. J Biochem 149, 131–143.

Tao, J., and Rolls, M.M. (2011). Dendrites have a rapid program of injury-induced degeneration that is molecularly distinct from developmental pruning. J Neurosci 31, 5398–5405.

Tung, T.T., Nagaosa, K., Fujita, Y., Kita, A., Mori, H., Okada, R., Nonaka, S., and Nakanishi, Y. (2013). Phosphatidylserine recognition and induction of apoptotic cell clearance by Drosophila engulfment receptor Draper. J Biochem 153, 483–491.

Vargas, M.E., Yamagishi, Y., Tessier-Lavigne, M., and Sagasti, A. (2015). Live Imaging of Calcium Dynamics during Axon Degeneration Reveals Two Functionally Distinct Phases of Calcium Influx. J Neurosci 35, 15026–15038.

Verdin, E. (2015). NAD(+) in aging, metabolism, and neurodegeneration. Science 350, 1208–1213.

Waller, A. (1850). Experiments on the Section of the Glossopharyngeal and Hypoglossal Nerves of the Frog, and Observations of the Alterations Produced Thereby in the Structure of Their Primitive Fibres. Philosophical Transactions of the Royal Society of London 140, 423–429.

Wang, J., Zhai, Q., Chen, Y., Lin, E., Gu, W., McBurney, M.W., and He, Z. (2005). A local mechanism mediates NAD-dependent protection of axon degeneration. J Cell Biol 170, 349–355.

Wen, Y., Parrish, J.Z., He, R., Zhai, R.G., and Kim, M.D. (2011). Nmnat exerts neuroprotective effects in dendrites and axons. Mol Cell Neurosci 48, 1–8.

Williams, P.R., Marincu, B.N., Sorbara, C.D., Mahler, C.F., Schumacher, A.M., Griesbeck, O., Kerschensteiner, M., and Misgeld, T. (2014). A recoverable state of axon injury persists for hours after spinal cord contusion in vivo. Nat Commun 5, 5683.

Witte, M.E., Schumacher, A.M., Mahler, C.F., Bewersdorf, J.P., Lehmitz, J., Scheiter, A., Sanchez, P., Williams, P.R., Griesbeck, O., Naumann, R., et al. (2019). Calcium Influx through Plasma-Membrane Nanoruptures Drives Axon Degeneration in a Model of Multiple Sclerosis. Neuron 101, 615–624 e615.

Xiong, X., Hao, Y., Sun, K., Li, J., Li, X., Mishra, B., Soppina, P., Wu, C., Hume, R.I., and Collins, C.A. (2012). The Highwire ubiquitin ligase promotes axonal degeneration by tuning levels of Nmnat protein. PLoS Biol 10, e1001440.

Yang, J., Wu, Z., Renier, N., Simon, D.J., Uryu, K., Park, D.S., Greer, P.A., Tournier, C., Davis, R.J., and Tessier-Lavigne, M. (2015). Pathological axonal death through a MAPK cascade that triggers a local energy deficit. Cell 160, 161–176.

Zhai, R.G., Cao, Y., Hiesinger, P.R., Zhou, Y., Mehta, S.Q., Schulze, K.L., Verstreken, P., and Bellen, H.J. (2006). Drosophila NMNAT maintains neural integrity independent of its NAD synthesis activity. PLoS Biol 4, e416.

Zhai, R.G., Rizzi, M., and Garavaglia, S. (2009). Nicotinamide/nicotinic acid mononucleotide adenylyltransferase, new insights into an ancient enzyme. Cell Mol Life Sci 66, 2805–2818.

